# Large-bodied ornithomimosaurs inhabited Appalachia during the Late Cretaceous of North America

**DOI:** 10.1101/2022.03.25.485782

**Authors:** Tsogtbaatar Chinzorig, Thomas Cullen, George Phillips, Richard Rolke, Lindsay E. Zanno

**Affiliations:** Paleontology Research Lab, North Carolina Museum of Natural Sciences, Raleigh, North Carolina, United States of America; Department of Biological Sciences, North Carolina State University, Raleigh, North Carolina, United States of America; Carleton University, Ottawa, Ontario, Canada; Conservation & Biodiversity Section, Mississippi Museum of Natural Science, Jackson, Mississippi, United States of America; Dow Chemical Company, Baton Rouge, Los Angeles, United States of America

**Keywords:** Eutaw Formation, Body-size evolution, Paleobiogeography, Ornithomimosauria, Theropod, Coexistence

## Abstract

Reconstructing the evolution, diversity, and paleobiogeography of North America’s Late Cretaceous dinosaur assemblages requires spatiotemporally contiguous data; however, there remains a spatial and temporal disparity in dinosaur data on the continent. The rarity of vertebrate-bearing sedimentary deposits representing Turonian–Santonian ecosystems, and the relatively sparse record of dinosaurs from the eastern portion of the continent, present persistent challenges for studies of North American dinosaur evolution. Here we describe an assemblage of ornithomimosaurian materials from the Santonian Eutaw Formation of Mississippi. Morphological data coupled with osteohistological growth markers suggest the presence of two taxa of different body sizes, including one of the largest ornithomimosaurians known worldwide. The regression predicts a femoral circumference and a body mass of the Eutaw individuals similar to or greater than that of large-bodied ornithomimosaurs, *Beishanlong grandis* and *Gallimimus bullatus*. The paleohistology of MMNS VP-6332 demonstrates that the individual was at least 11 years of age (similar to *B. grandis* [∼375 kg, 13–14 years old at death]). Additional pedal elements share some intriguing features with ornithomimosaurs yet suggest a larger-body size closer to *Deinocheirus mirificus*. The presence of a large-bodied ornithomimosaur in this region during this time is consistent with the relatively recent discoveries of early-diverging, large-bodied ornithomimosaurs from mid-Cretaceous strata of Laurasia (*Arkansaurus fridayi* and *B. grandis*). The smaller Eutaw taxon is represented by a tibia preserving seven growth cycles, with osteohistological indicators of decreasing growth, yet belongs to an individual with near reaching somatic maturity of the larger taxon, suggesting the co-existence of medium- and large-bodied ornithomimosaur taxa during the Late Cretaceous Santonian of North America. The Eutaw ornithomimosaur materials provide key information on the diversity and distribution of North American ornithomimosaurs and Appalachian dinosaurs and fit with broader evidence of multiple cohabiting species of ornithomimosaurian dinosaurs in Late Cretaceous ecosystems of Laurasia.

## Introduction

During the majority of the Late Cretaceous, the southern North American continent was divided into two landmasses by the expansion of the Western Interior Seaway, forming Laramidia to the west and Appalachia to the east (e.g., [1–3]). Continental separation had appreciable consequences for the evolution of North American dinosaurs, with distinct lineages evolving in isolation on each landmass [4]. Although the vertebrate fossil record of Appalachia suggests a distinct and diverse fauna [5,6], the majority of this record is based on relatively poorly preserved, and often isolated specimens, when compared to the more extensive record of Laramidian taxa [7,8]. This is due, in part, to preservational and collection biases, as the vast majority of the exposed sedimentary units in Appalachia represent marine deposits [9,10] and preserved fossils of terrestrial taxa are often fragmentary [5,6]. Thus, the dinosaur fossil record of Appalachia is poor when compared to the extensive record of terrestrial fluvial and coastal plain deposits in Laramidia. Nonetheless, knowledge of the Mesozoic terrestrial fauna of eastern North America is rapidly growing. From these often-isolated elements, researchers have been able to piece together a diverse Appalachian vertebrate fauna represented by hadrosauroids, ceratopsids, and theropods [11–17]. These discoveries have greatly strengthened our understanding of the evolution, biodiversity, and paleoecology of the Appalachian dinosaur fauna [3,10,11,13,17–22]. Yet much remains to be learned. One group with particularly poor representation is maniraptoriform theropods. Maniraptoriforms represented in Appalachian assemblages include dromaeosaurids and ornithomimosaurs (e.g., [15,22,23]).

Ornithomimosaurs are lightly built theropod dinosaurs characterized by long fore limbs, powerful hind limbs and relatively small skulls that exhibit reduced teeth to fully edentulous beaks [24–26]. Although the clade is well represented in the Upper Cretaceous deposits (Campanian-Maastrichtian) of the Western Interior of North America [24,25], the fossil record is relatively rare in more older eastern North American localities [6,26,27]. A recent review of the fossil record from this subcontinent cited several ornithomimosaur specimens described during the last decade; however, most of these specimens are too fragmentary to be diagnosed to species level [23,28]. These include associated vertebrae and isolated elements of the hind limb from the Lower Cretaceous Arundel Clay (Potomac Formation) of Maryland [15,29], which are undiagnostic at finer taxonomic levels [30]. Recent new description of an early-diverging ornithomimosaur taxon, *Arkansaurus fridayi*, from Arkansas [14] represents an exception to this pattern; however, *A. fridayi* predates the isolation of Appalachia and Laramidia and cannot therefore inform us on the impact of isolation on the evolution of eastern North American dinosaurs. Moreover, the near absence of body fossil records of ornithomimosaur dinosaurs across the whole of North America (including both Laramidia and Appalachia) continent from the Cenomanian to the Campanian (a gap of ∼10 Myrs.), is currently obscuring the macroevolutionary history and paleobiogeography of ornithomimosaurs more broadly on this continent.

Here, we describe multiple specimens of ornithomimosaurian dinosaur from the Santonian Eutaw Formation, assembled by the Mississippi Museum of Natural Science, from Lowndes County, Mississippi, in a limited exposure along Luxapallila Creek (Fig 1). The Eutaw ornithomimosaur materials represent individuals of different body sizes, therefore we test for the presence of multiple taxa using osteohistological interpretations. We then discuss the implications of these specimens on our understanding of ornithomimosaur body size evolution and diversity.

**Fig 1.**
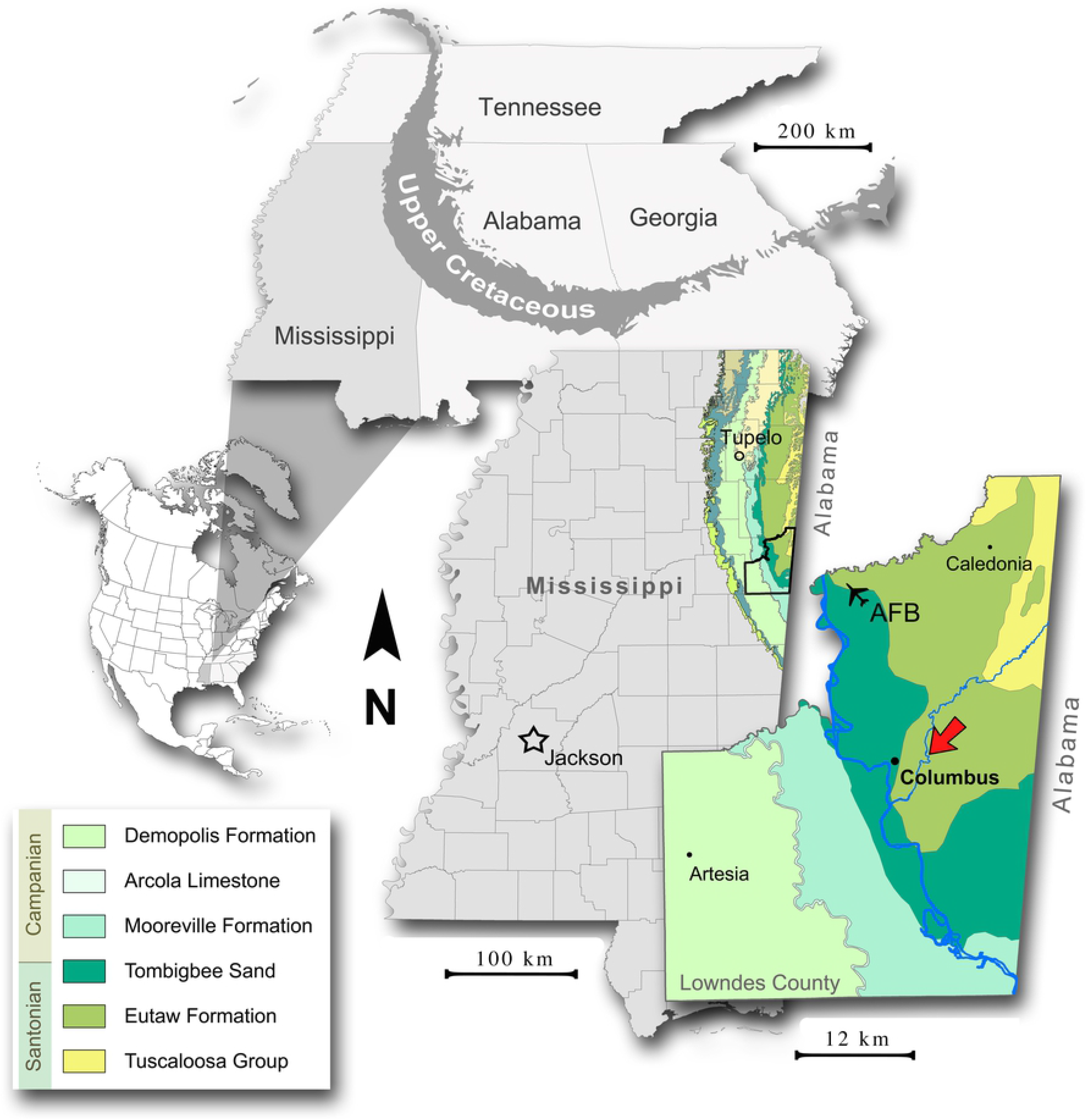
Geographic and stratigraphic occurrence of the Eutaw ornithomimosaurs’ assemblage. Geologic map of Lowndes County with red arrow showing location of fossil site. A simplified geologic key is provided with placement of the Santonian-Campanian boundary in Mississippi based on Dowsett [31], Kennedy & Cobban [32], Mancini et al. [33], and Puckett [34].

## Materials and Methods

### Specimens

**The elements referable to large-bodied ornithomimosaurs** - Incomplete astragalus (MMNS VP-8826); a nearly complete, pathologic second metatarsal (MMNS VP-6332, MT-II); distal halves of the third (MSC 13139, MT-III) and the fourth (MMNS VP-6183, MT-IV) metatarsals; pedal phalanges MMNS VP-4955 (PII-1), 9444 (PII-1), 4949 (PIII-1), and 7119 (PIV-2).

**The elements referable to medium-bodied ornithomimosaurs -** Partial dorsal centra (MMNS VP-113 and MMNS VP-6120); incomplete posterior caudal centrum (MMNS VP-6329); complete manual phalanx (MMNS VP-6419, PIII-1); complete manual ungual (MMNS VP-2963); and incomplete tibial shaft (MMNS VP-7649). These elements are described and figured in the supplementary information.

### Osteohistological thin-sectioning

A total of three osteohistological thin-sections were made from the tibia (MMNS VP-7649) and the second metatarsal (MMNS VP-6332). These elements represent two different individuals based on relative size. The sections of MMNS VP-6332 were taken from the proximal- and the mid-shaft, whereas only the mid-shaft of MMNS VP-7649 was sectioned from the specimen. Prior to consumptive sampling, thin-sectioned specimens were photographed, measured, molded, and cast. These thin-sections were cut using a Ryobi tile saw (model WS750L) before embedded and affixed to plexiglass slides using a clear epoxy resin (EPO-TEK 301) and cut with a Buehler Isomet 1000 diamond wafer blade, low-speed precision saw. The grinding/polishing processes (down to a thickness of approximately 50-60 μm) were done on a Hillquist thin-section machine and Buehler Metaserv-250 grinder and polishing machine using a series of abrasive paper disks with increasing grit sizes (400, 800, 1200), and a microcloth. All prepared thin-sections were examined using a Nikon Eclipse Ci Pol polarizing microscope and photographed with a Keyence VHX-5000 microscope. All thin-sections were made in the histological facility of the Paleontological Research Laboratory, North Carolina Museum of Natural Sciences, using standard paleo-osteohistological methods [35].

### Body mass estimates

To estimate the body masses of two of the largest individuals (MMNS VP-6332 and MMNS VP-7119), and the one smaller individual (MMNS VP-7649) from our sample, we performed ordinary least squares (OLS) linear regressions to predict a femoral circumference (FC) from original measurements of ornithomimosaurian elements (FC – pedal phalanx IV-2 length, FC - metatarsal II length, and FC - tibia length). The bauplan of *Deinocheirus mirificus* is unusual among ornithomimosaurs. Skeletal ratios of *D. mirificus* are outliers in our dataset and had a pronounced effect on estimated FC. Given also that the phylogenetic relationships of the Eutaw taxa are unknown, we chose to predict a range of FC (and therefore masses) for MMNS VP-6332, MMNS VP-7119, and MMNS VP-7649 using datasets that both include and exclude *D. mirificus*. All measurement and estimation data related to these analyses are presented in S1 Table, with regression results displayed in S1 Figure. We used the R package ‘MASSTIMATE” [36], based on extant scaling relationships between FC and BM and modified for use with bipeds (cQE; Campione et al. [43]), to estimate body mass (BM) from FC. Masses of smaller/juvenile individuals were estimated by applying the ‘developmental mass extrapolation’ (DME) approach of Erickson and Tumanova [38] to the estimated femoral circumference values of smaller specimens and the femoral circumference and body mass values of large/adult specimens (*sensu* Chiba 2018) (S2 Table).

### Geological Setting & Locality Information

The Upper Cretaceous strata of the Eutaw Group are exposed in northeastern Mississippi as part of an outcrop belt that extends from west-central Georgia, through central Alabama, and into western Tennessee by way of northeastern Mississippi [39] (Fig 1). Within Mississippi and western Alabama, the Eutaw Group, in ascending superposition, comprises the McShan Formation, the lower, unnamed member of the Eutaw Formation, and the Tombigbee Sand [40] (Fig 2A). The contact of the Tombigbee Sand with the lower Eutaw is represented by an erosional unconformity and begins a major transgressive depositional sequence with the former succeeded by the Mooreville Formation [33,39,40]. Although the Tombigbee is very distinct from the lower Eutaw in both lithologic and sequence stratigraphic terms [33,40,41], it has long been relegated to a member of the Eutaw Formation by most stratigraphic works. The Tombigbee-Mooreville contact is diachronous, dating to the earliest Campanian in east-central Mississippi and Santonian eastward into central Alabama [31,32] (Fig 2A). Biostratigraphic and radiometric analyses have yielded ages of late Santonian through earliest Campanian for the Tombigbee Sand in north Mississippi (e.g., [32,33,42]). The lower Eutaw Formation falls within the *Dicarinella concavata* interval zone, and the lowest portions of this unit extend into the Coniacian [34]. The lower Eutaw is interpreted as a complex of regressive facies immediately preceding the Tombigbee-Mooreville transgressive beds [33].

**Fig 2.**
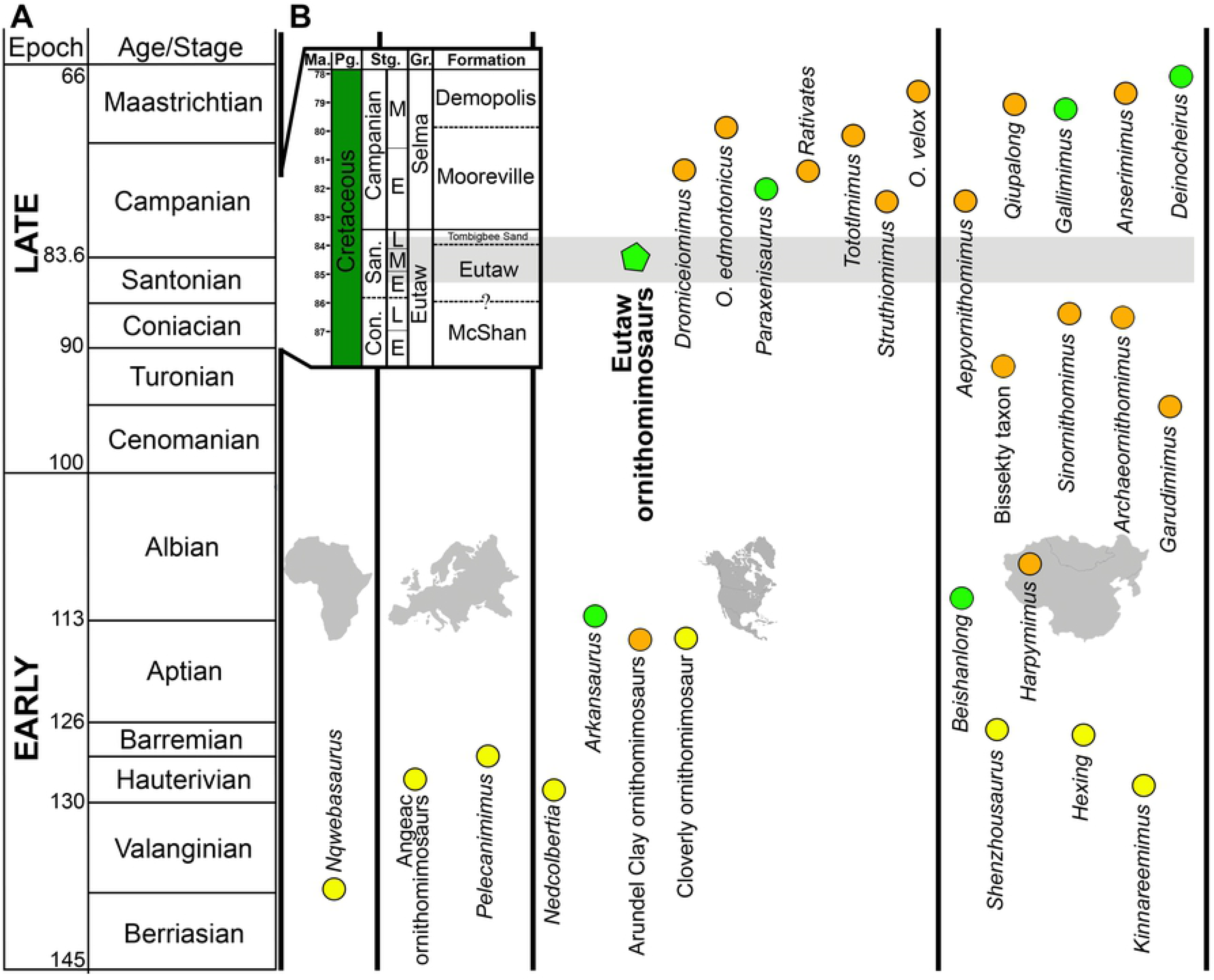
Temporally calibrated chart of Ornithomimosauria, showing the continental distribution of the known taxa and binned body mass. (**A**) General stratigraphic column of the Cretaceous Period, showing the global distribution of ornithomimosaur taxa divided by continents (inset stratigraphic column shows the local stratigraphy of Mississippi County, where the Eutaw ornithomimosaurs were discovered); (**B**) The Eutaw ornithomimosaurian taxa are indicated by a green pentagon. Colored circles refer to the different body mass of ornithomimosaur taxa: yellow circles indicate small-bodied ornithomimosaurs (3.58 kg - 34.78 kg), orange circles indicate medium-bodied ornithomimosaurs (60.76 kg – 233.83 kg), and green circles indicate large-bodied ornithomimosaurs (>380.48 kg). The figure is modified from Hunt and Quinn [14].

All specimens described herein were recovered from two locations—MS.44.001a (to the south) and MS.44.001b (to the north)—1.3 km apart in the same stratigraphic interval within the upper part of the lower Eutaw Formation exposed along a channelized length of Luxapallila Creek in Columbus, Lowndes County, Mississippi (Fig 1). The macrofossil-bearing bed within the Luxapallila section (MS.44.001) is a relatively thin (typically 20-90 cm thick) condensed zone of fine-sandy clay containing a variety of coarse components at its base and fining upwards. Composition of the basal lag, coined the “Lux lag” [43] includes bones, teeth, vertebrate coprolites, moldic invertebrates, lignitized wood, phosphatized wood (reworked), sundry phosphatized rounded and frequently flattened pebbles, rare exotic clasts, and regularly occurring claystone rip-ups from the indurated surface on which it rests. Along the exposed section, this macrofossiliferous condensed zone is typically bounded above and below by thinly laminated, occasionally rippled, lignitic clays, although immediately subjacent to the fossil bed at MS.44.001b is a suite of cross-bedded and flaser-bedded clean sands with lignitic interlaminae and regularly occurring burrows (including *Ophiomorpha*). These bounding beds are typical of the lower Eutaw [39,44,45] and, to some extent, the McShan [46,47]. Although the exposed section of MS.44.001 has not yielded useful standardized index fossils, like ammonites and nannofossils, the late Santonian age of the vertebrate-bearing interval is reasonably straightforward from both its relative stratigraphic position below (≥ 10 m) the typical Tombigbee lithology to the south and west [39,44] and ichthyofaunal species composition [43,48,49].

A similar described section to that bearing the Lux lag occurs almost 30 km to the north along strike, where a vertebrate-rich bed in the upper part of the lower Eutaw lies well below the typical Tombigbee Sand [45]. Even further northward along strike, some workers have ascribed a dinosaur-bearing lithofacies to the subjacent McShan Formation [1,42]. Some workers have ascribed a dinosaur-bearing lithofacies, even further northward along strike, to the McShan Formation [1,42], but the ephemeral exposure was not available for detailed study and is interpreted here as stratigraphically equivalent to the lower Eutaw in the Columbus area. To the east and northeast, marginal marine lithofacies attributable to the McShan have been reported by Cook [46], although they notably lack burrowing, the presence of which is characteristic of the Tombigbee Sand to the south and west of MS.44.001 but also the aforementioned sand beds in the vicinity of the upstream fossil locality. The McShan Formation was erected by Monroe et al. [47], who separated and distinguished it from the superjacent Eutaw by a regional unconformity with a pebbly bed containing shark teeth in the basal portion of the latter. Although this coarse clastic basal Eutaw bed was studied primarily in western Alabama [47], it is readily correlate to the Lux lag. As the McShan Formation is often indistinguishable in subsurface analyses, its usage has been abandoned by many workers, who treat the unit as part of the lower Eutaw [33,50].

Vertebrate fossils from the Lux lag at MS.44.001 were first documented by Kaye [44] who reported shark teeth only and also described the interval as below the typical Tombigbee Sand or within a “transitional zone”. The general diversity of vertebrate remains was first reported, without details, by Phillips & Loftis [51], but a selachian assemblage and lungfish tooth plate were described at length by Cicimurri et al. [43] and Harrell & Ehret [52], respectively. The vertebrate species represented at this site consist of an ecologically disparate mix of various marine, brackish, freshwater, and even terrestrial taxa [43,52]. Typical of lag concentrations, skeletal dissociation is high. Marine fish remains are the most common vertebrate constituents of the Lux lag, whereas crocodilian and chelonian remains occur with lesser frequency. Rarer still are the remains of plesiosaur, mosasaur, and dinosaur, although the incidence of dinosaur bone (mostly fragmentary) outnumbers that of the marine reptiles. The dinosaurian assemblage from MS.44.001 is perhaps the most significant component of the fauna because it is richer in total individual remains than any other Mississippi site, particularly given the limited size of the exposure. The Lux lag contains a diverse dinosaurian assemblage, at least for marginal marine facies; it includes the fragmentary remains of hadrosaur, nodosaur, and a variety of theropods. **Taphonomy** - The thin, regionally extensive Lux lag is a temporally and environmentally constrained interval within the regressive (lowstand) beds of the lower Eutaw Formation [33]. Monroe et al. [47] erected the McShan, in part, due to a coarse, fossiliferous facies he identified regionally in the base of the lower Eutaw (sensu stricto). Although Mancini et al. [33] did not incorporate this facies change at the McShan-Eutaw contact in their regional sequence stratigraphic model, they seemed to confine it to a relatively narrow temporal interval (Santonian) and depositional environment (estuarine). This point is crucial to the discussion about the relative coexistence of component taxa, especially the theropod dinosaurs—the subject of this paper. The dinosaurian elements, as is the case with basically all the observed phosphatized vertebrate clasts within the Lux lag, are inseparable as to related degrees of surface erosion. Large fragmentary bone dominates the dinosaurian component of the assemblage, but the gradual accumulation of more complete elements over ∼25 years (following stream channelization) have made the current project possible.

As previously discussed, close proximity of the superjacent, well-dated Tombigbee Sand allows a minimum age of late Santonian for the Lux lag. The lack of a bone-bearing subjacent facies as a potential clast-contributor to the Lux lag suggests temporal mixing is minimal to nonexistent, unless of course any vertebrate-bearing facies were removed earlier in the lowstand. Any reworking of older clast constituents from such hypothetical earlier Eutaw fossil beds could have introduced Coniacian (or older) elements. However, the assessment of the selachian assemblage [43] produced no elements that were strictly pre-Santonian, and the marine faunal character was essentially identical to that of better dated marine vertebrate assemblages nearby [48]. In addition, no pre-Santonian bone beds have been identified in the Upper Cretaceous of the region.

Formation of the Lux lowstand lag is beyond the scope of this study, but its ecological content is very revealing. The component taxa represent a mixture of paralic and very shallow marine vertebrate components. The proximity of fluvio-riparian habitats is suggested by the dinosaurs, lungfish [52], and kinosternoid turtles (work in progress); estuarine habitats by *Atractosteus*, crocodilian, and hybodont shark remains; and marine origins for the remainder of the aquatic taxa, mostly fishes. However, deep-water marine taxa are notably rare. Pelagic forms, like mosasaurs, plesiosaurs, and the ginsu shark *Cretoxyrhina* are exceptionally scarce occurrences. Several of the selachian taxa that are otherwise also common in deeper neritic environments are frequently represented in the Lux lag by juvenile remains, as in the relative abundance of early ontogenetic stages of goblin shark (*Scapanorhynchus*) and myliobatid ray (*Brachyrhizodus*) teeth [43]. Most selachians are born and raised in coastal marshes and estuaries [53], thus their relative abundance in the Lux lag suggests an equivalent depositional environment. The sedimentology of the lower Eutaw Group (i.e. McShan) further supports an estuarine complex [33,46]. Thus, although the geographic scope of this mixed assemblage is seemingly diverse, the assemblage is actually confined to area of overlapping coastal ecosystems and within a relatively narrow chronostratigraphic interval, making a compelling case for the coexistence of its dinosaurian elements.

## Institutional Abbreviations

**AM**, Albany Museum, Grahamstown, South Africa; **AMNH**, American Museum of Natural History, New York City, NY, USA; **ANSP**, Academy of Natural Sciences, Philadelphia, PA, USA; **BENC**, Benemérita Escuela Normal de Coahuila, Mexico; **BMRP**, Burpee Museum of Natural History, Rockford, IL, USA; **CMN,** Canadian Museum of Nature, Ottawa, Ontario, Canada; **FMNH PR**, Field Museum of Natural History, Chicago, IL, USA; **FRDC-GJ**, Fossil Research and Development Center, Third Geology and Mineral Resources Exploration Academy, Gansu Provincial Bureau of Geo-Exploration and Mineral Development, Lanzhou, China; **HGM**, Henan Geological Museum, Henan Province, China; **IVPP**, Institute of Vertebrate Paleontology and Paleoanthropology, Beijing, China; **LACM**, Los Angeles County Museum, LA, USA; **LH**, Long Hao Institute of Geology and Paleontology, Hohhot, China; **MMNS VP**, Mississippi Museum of Natural Science, MS, USA; **MOR**, Museum of the Rockies, Bozeman, MT, USA; **MPC-D**, Institute of Paleontology, Mongolian Academy of Sciences, Ulaanbaatar, Mongolia; **MSC**, McWane Science Center, Birmingham, AL, USA; **NCSM**, North Carolina Museum of Natural Sciences, Raleigh, NC, USA; **ROM**, Royal Ontario Museum, Toronto, ON, Canada; **TMP**, Royal Tyrrell Museum of Paleontology, Drumheller, AB, Canada; **RMM**, Red Mountain Museum, Birmingham, AL, USA; **SMU**, Southern Methodist University, Dallas, TX, USA; **YPM**, Yale Peabody Museum of Natural History, New Haven, CT, USA; **UAM**, University of Arkansas Museum, University of Arkansas, Fayetteville, AR, USA; **UCMZ (VP)**, University of Calgary Museum of Zoology, Calgary, Alberta, Canada; UMNH VP, Utah Museum of Natural History, Salt Lake City, UT, USA; **ZIN PH**, Paleo-herpetological Collection, Zoological Institute, Russian Academy of Sciences, Saint Petersburg, Russia; **ZPAL**, Institute of Paleobiology, Polish Academy of Sciences, Warsaw, Poland.

## Results

### Systematic palaeontology

Dinosauria Owen, 1842 [54]

Theropoda Marsh, 1881 [55]

Ornithomimosauria Barsbold, 1976 [56]

Ornithomimosauria indet.

### Description

#### Astragalus

An incomplete left astragalus (MMNS VP-8826) is preserved, missing its medial condyle and most of the ascending process, which is represented only by its base on the lateral side (Fig 3). The preserved portion of the lateral condyle is spherical. Although the medial condyle is missing, enough anatomical information is present to determine in MMNS VP-8826 that the lateral condyle is more extensive anteriorly and the two condyles are separated by a well-developed intercondylar bridge (Fig 3E). The bridge of MMNS VP-8826 is not strongly developed as those of tyrannosauroids (e.g., *Dryptosaurus aquilunguis*) and troodontids (e.g., *Talos sampsoni* and *Gobivenator mongoliensis*), which exhibit highly constricted round or V-shaped outlines of the intercondylar bridge between the lateral and medial condyles in proximal view (S2B and S2F Fig) [57,58]. The base of the ascending process of MMNS VP-8826 bears a shallow median fossa (Fig 3A, D). A shallow median fossa is unlike those of deeper median fossae of the tyrannosauroids *Appalachiosaurus montgomeriensis*, *D. aquilunguis,* and *Tyrannosaurus rex* (S2A1-C1 Fig), as well as the caenagnathoid *Caenagnathus collinsi* and the therizinosauroid *Falcarius utahensis* (S2D1-E1 Fig), but it resembles those of ornithomimosaurs (S2G1-I1 Fig). The horizontal groove is commonly present at the base of the ascending processes in some theropods, such as ornithomimosaurs, tyrannosauroids, and troodontids (e.g., *T. sampsoni*), but it is completely absent in therizinosauroids (e.g., *F. utahensis*) [57,59], and the troodontids *G. mongoliensis* and *Sinornithoides youngi* [58]. A well-pronounced horizontal groove is partially preserved at the base of the preserved ascending process of MMNS VP-8826 (Fig 3A, D). This groove is much shallower than those of the tyrannosauroid *A. montgomeriensis* and the troodontid *T. sampsoni* (S2F1 Fig) [11,57]. The contact surfaces for the articulation of the anterior and distal surfaces of the tibia are flat and straight in lateral and medial views like most theropod dinosaurs (Fig 3B, D). The base of the astragalar body is slightly constricted distally and nearly straight for the tibia’s distal articulation in posterior view (Fig 3A-B). When articulated with the tibia, the distal end of the astragalar body would seen in posterior view as in ornithomimosaurs such as *Aepyornithomimus tugrikinensis, Ornithomimus velox,* and *Qiupalong henanensis* [60–62]. The portion of the astragalar body underlying the tibia (astragalar base) is dorsoventrally thick and laterally unflared of MMNS VP-8826 is consistent with the condition of ornithomimosaurs (Fig 3B-D) and contrasts with that of tyrannosaurs such as *T. rex* (e.g., MOR 1125 and FMNH PR 2081) [63], and *A. montgomeriensis* [11] (S2A2-C2 Fig).

**Fig 3.**
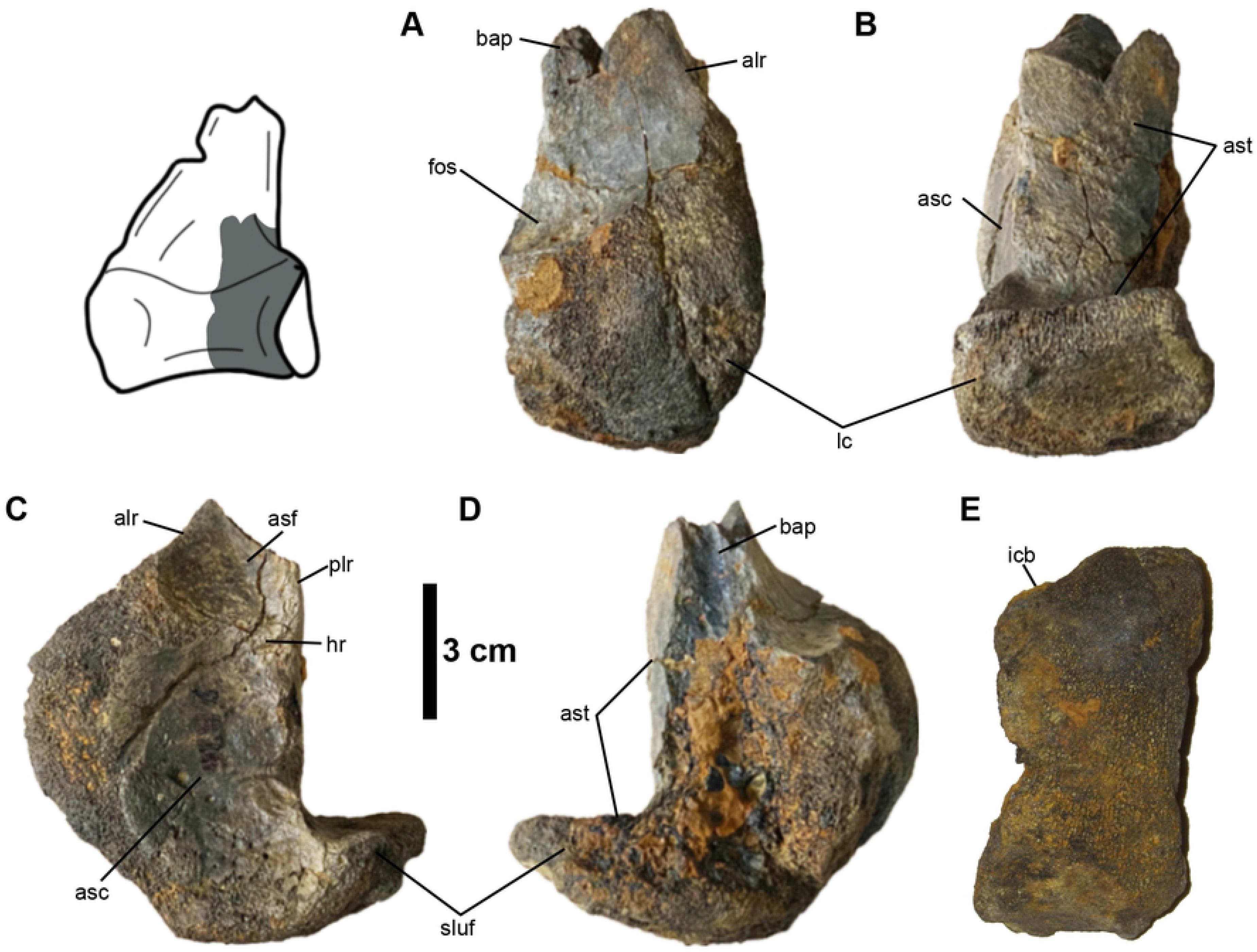
Left astragalus (MMNS VP-8826). **(A)**, anterior; **(B)**, posterior; **(C)**, lateral; **(D)**, medial; **(E)**, distal views. Interpretive illustration of *Q. henanensis* (HGM 41HIII-0106) shows the approximate location of the preserved portion of the astragalus. Abbreviations: alr, anterolateral ridge; asc, articular surface for the calcaneum; asf, articular surface for the fibula; ast, articular surface for the tibia; bap, base of the ascending process; fos, median fossa; hr, horizontal ridge; icb, intercondylar bridge; lc, lateral condyle; plr, posterolateral ridge; sluf, laterally unflared articular surface.

In MMNS VP-8826, the well-preserved articular surfaces of MMNS VP-8826 for the fibula and the calcaneum are smooth and concave and are separated by a weak horizontal ridge (Fig 3C). The fibular articular surface is restricted by prominent ridges anteriorly (anterolateral ridge) and posteriorly (posterolateral ridge), forming a posterolaterally facing anterior surface and anterolaterally facing posterior surface (Fig 3C). In anterior view, the anterolateral ridge is relatively vertically oriented (Fig 3A), compared to the ornithomimid *Q. henanensis* as well as those of tyrannosauroids (e.g., *A. montgomeriensis* and *T. rex* [MOR 1125]) (S2A1, S2C1, and S2G1 Fig), which slope medially forming a concave margin. Unlike the astragali of *Anzu* (NCSM 33801), *T. rex* (MOR 1125), and *F. utahensis* (UMNH VP 12364) (S2A2, D2, and E2 Fig), there is no deep junction between the base of the ascending process and the lateral condyle of MMNS VP-8826 in lateral view (Fig 3C). MMNS VP-8826 lacks a well-developed notch centered on the anterolateral margin of the lateral condyle that is present on some ornithomimids (e.g., *G. bullatus, Q. henanensis* and large Gansu ornithomimid [46,50,51]), and to a lesser degree troodontids (e.g., *T. sampsoni*) and therizinosauroids (e.g., *F. utahensis*) (S2D1 and S2F1 Fig). In this respect, the condition of MMNS VP-8826 is similar to the ornithomimid *A. tugrikinensis* and caenagnathids like *Anzu wyliei* and *Gigantoraptor erlianensis* [66,67]. This notch is different from the notch on the anterolateral margin of the lateral condyle that is described in tyrannosauroids. Whereas the notch of the tyrannosauroid *D. aquilunguis* is more distally located on the lateral condyle, *A. montgomeriensis* exhibits a proximally located notch, which is absent on MMNS VP-8826 [11,68] (S2B3 and S2C2 Fig).

The border of the articular surface of calcaneum is convex anteriorly and straight anterodistally in lateral view (Fig 3C), similar in form to the tyrannosauroids *A. montgomeriensis*, *D. aquilunguis* (S2B2-C2 Fig), and some ornithomimosaurs, such as *A. tugrikinensis* and *Q. henanensis* (S2G2-H2 Fig). However, it is different from caenagnathoids, therizinosauroids, troodontids, and tyrannosaurids. Whereas those of *T. rex* (MOR 1125) and *F. utahensis* exhibit a prominent sulcus anteriorly on the anterolateral margin of the astragalar body in relation to the horizontal groove on the anterior surface (S2A2 and S2D2 Fig), large-bodied caenagnathids, such as *C. collinsi* from the Dinosaur Park Formation and *Anzu* sp. (NCSM 33801), as well as the troodontid *T. sampsoni,* display a round outline of the calcaneal articular surface in lateral view (S2E2-F2 Fig) [57,69]. Moreover, the dorsoventrally thick and laterally unflared articular surface of the astragalar base of MMNS VP-8826 is not referable to the aforementioned theropod groups, such as the tyrannosauroids *A. montgomeriensis, D. aquilunguis*, and *T. rex*, and therizinosauroid *D. utahensis* as well as *C. collinsi* (S2A2-D2 Fig). Furthermore, MMNS VP-8826 is also differentiated from these theropod groups by the absence of a deep juncture between the anterolateral ridge and the lateral condyle (S2A2 and S2D2-E2 Fig).

In short, the visibility of the astragalar body in posterior view, presence of a less constricted intercondylar bridge, shallow horizontal groove, absence of a laterally flaring astragalar base, well-developed notch on the anterolateral margin of the lateral condyle, and more vertically oriented anterior ridge of the fibular contact allows us to support an ornithomimosaur affinity and refute a tyrannosaur referral.

#### Metatarsus

Three metatarsals are represented in the Eutaw ornithomimosaur assemblage, including the second (MMNS VP-6332), third (MSC 13139), and fourth (MMNS VP-6183) metatarsals. These appear to belong to different individuals (Figs 4-6).

**Fig 4.**
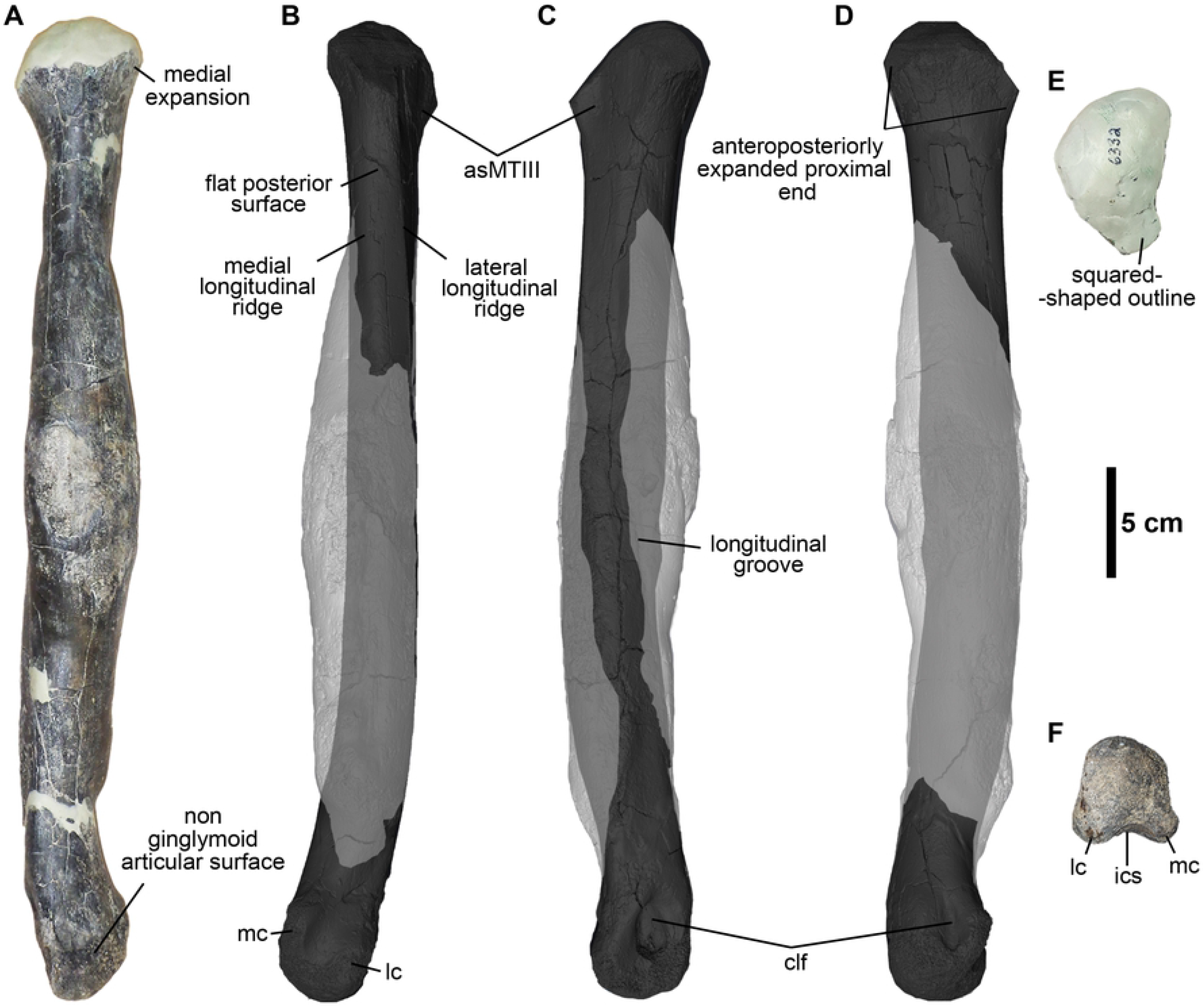
Right second metatarsal (MMNS VP-6332). **(A)**, anterior; **(B)**, posterior; **(C)**, lateral; **(D)**, medial; **(E)**, proximal; **(F)**, distal views. Abbreviations: asMTIII, articular surface for the third metatarsal; clf, collateral ligament fossa; ics, intercondylar sulcus; lc, lateral condyle; mc, medial condyle. (**A, E, F**) gross morphology of original bone; (**B-D**), 3D model of the metatarsal shows the original morphology of the shaft in dark gray and distribution of a pathology in light gray.

#### The second metatarsal (MT-II)

MMNS VP-6332 is essentially complete yet missing the proximal-most articular surface and most of the shaft is deformed due to a pathology (Fig 4A). MMNS VP-6332 is identified as a second metatarsal of the right foot based on the subtriangular proximal end with a nearly flat articular surface for the articulation of the third metatarsal, medially deviated distal end with a transversely unconstricted, quadrangular distal articular surface relative to its height, and a less flared medial condyle (Fig 4). Although MMNS VP-6332 is not complete, we used the length of the preserved portion of the metatarsal to estimate the length of the complete element at ∼434 mm. This estimate is similar in size to the second metatarsal of the large Gansu ornithomimid from China [65], ∼17% longer than those of the Early Cretaceous ornithomimosaurs *A. fridayi* and *Beishanlong grandis* [14,70], and more than twice the length of the geographically closest type specimen *O. velox* [61].

MMNS VP-6332 is proximodistally long and slender and the proximal two-thirds of the shaft is straight, with only ∼20% of the distal articular caput deviating medially, similar to most ornithomimosaurs (Fig 4A). The buttressing surface, which is located on the lateral surface of the distal half of the second metatarsal, is one of the differentiating characteristics between ornithomimosaurs and tyrannosauroids (e.g., [57–59]). Based on architecture of the non-pathological cortical bone periosteal surface segmented via CT reconstruction (Fig 4B-D), there is no evidence of the buttressing surface on MMNS VP-6332, indicating that this metatarsal is not referable to tyrannosauroids.

Proximally, MMNS VP-6332 is expanded anteroposteriorly and medially relative to the shaft as in most theropods (Fig 4A and D). However, this expansion is less than that of tyrannosauroids (S3 Fig) but closely resembles the condition of ornithomimosaurs. The shaft just distal to the proximal end is not affected by pathology, preserving the original morphology of the shaft on gross inspection (Fig 4). The morphology of the proximal shaft is typical of ornithomimosaurs, with a suboval cross-section (wider anteroposteriorly than mediolaterally), displaying flat lateral and posterior surfaces, and convex medial and anterior surfaces (Fig 4B) like *Dromiceiomimus brevitertius* and *Rativates evadens*, which also exhibit a similar flat surface on the posterior surface of the second metatarsal [74,75]. But it differs from the late-diverging ornithomimosaur taxa, such as *O. velox*, *G. bullatus*, and *Struthiomimus altus* as well as those of early-diverging ornithomimosaurs *A. fridayi, B. grandis* and *Harpymimus okladnikovi*, which have a relatively convex posterior surface. Overall, the aspect of the proximal end indicates that MMNS VP-6332 is referable to a taxon with an arctometatarsalian foot condition [25,65], which is similar in morphology to the type specimen of *O. velox*, and a large Gansu ornithomimid [61,65,76]. Although both ornithomimosaurs and tyrannosaurs exhibit an arctometatarsalian foot, the tyrannosauroids *A. montgomeriensis*, *Bistahieversor sealeyi*, *Gorgosaurus libratus*, *Tarbosaurus bataar,* and *T. rex* as well as Delaware tyrannosauroid metatarsal, exhibit a deeply notched articular surface for the third metatarsal on the corresponding surface of the second metatarsal [11,63,77]; whereas the articular surface for MT-III on MMNS VP-6332 is relatively flat and weakly notched as in ornithomimosaurs [62,64,78] (Fig 4E).

The proximal and distal ends are slightly rotated clockwise in relation to the main axis, resulting in a weakly twisted metatarsal shaft (Fig 4). Although this feature is unusual in ornithomimosaurs, the medially rotated distal articular end is also reported in metatarsals of the large Gansu ornithomimid, in which the articular surface is slightly inclined distally and medially [65]. Although the degree of rotation in MMNS VP-6332 is greater than that observed in the large Gansu ornithomimid, it is possible that this may relate to pathological deformation.

The posterior surface of the pathologically unaffected proximal shaft of MMNS VP-6332 bears a flat surface (mediolaterally ∼1 cm wide), which is bordered by prominent longitudinal ridges laterally and medially as in *O. velox* and the large Gansu ornithomimid (Fig 4B) [61,65]. The lateral ridge is more extensive proximodistally than the medial ridge, extending to the pathologically affected bone and then obscured due to pathology. The flat surface has a square-shaped outline in cross-sectionally, distinguishing it from caenagnathoids, such as *C. collinsi* and other referred specimens (e.g., TMP 1993.036.0197 and TMP 1993.036.0198), which exhibit a convex surface [69,79]. Furthermore, the prominent longitudinal groove exists along the anterolateral surface of MMNS VP-6332 in lateral view (Fig 4C). Although this groove is relocated from the original position due to pathological deformation, it is apparent that it represents the anterior border of the articular surface for the MT-III.

The articulation for the metatarsal-phalangeal joint of the distal articular surface is non-ginglymoid; rather, it has a smooth and bulbous articular surface as in other ornithomimosaurs (Fig 4A and F). Transversely, the distal articular caput is slightly broader than the width of the shaft in anterior view like *O. velox* and the large Gansu ornithomimid (Fig 4F). The height/width ratio of the distal articular caput is subequal, with equally developed two distal condyles, forming a round bulbous shape in posterior view. The condyles have straight lateral and slightly concave medial outlines and are separated by a relatively deep, broad intercondylar sulcus in distal view. The lateral condyle is slightly larger than the medial condyle, particularly on the posterior surface. The bulbous medial condyle of MMNS VP-6332 is unlike that of the narrow and more sharply ridged medial condyles of late-diverging ornithomimosaurs, such as *O. velox*, *A. tugrikinensis,* and the large Gansu ornithomimid, as well as early-diverging ornithomimosaurs (e.g., *H. okladnikovi,* [66]). It exhibits shallower intercondylar sulcus than observed on *O. velox*. Furthermore, it bears well-developed collateral ligament fossae (Fig 4C-D). The shape of these collateral ligament fossae is equally ellipsoid, but the lateral collateral ligament fossa is proportionately larger and deeper than the medial one. They are positioned at approximately the center of the distal caput. The shape of the collateral ligament fossae is different from that of *O. velox*, which bears round ligament fossae in outline [61].

Broadly speaking, theropods bearing arctometatarsalian feet include deinonychosaurs, caenagnathoid oviraptorosaurs, ornithomimosaurs, and tyrannosauroids. However, the presence of the non-ginglymoid distal articular surface of MMNS VP-6332 differentiates this element from that of deinonychosaurs (e.g., *Adasaurus mongoliensis*, *D. antirrhopus, Dromaeosaurus albertensis,* and *Velociraptor mongoliensis*) [81–84]. A square-shaped outline of the posterior border of the proximal articular surface, lack of the posteromedial ridge along the shaft, and the convex, non-ginglymoid distal articular surface with a posteriorly blunt lateral and medially more deflected medial condyles rules out caenagnathoid oviraptorosaurs [69,79]. In general, the combination of a straight, slender shaft, expanded proximal end, and non-ginglymoid distal articular surface of MMNS VP-6332 broadly resembles those of the Appalachian tyrannosauroids, such as *A. montgomeriensis* and the Delaware tyrannosauroid (S3 Fig), as well as those of slender footed tyrannosauroids, including *Alectrosaurus olseni* and *Moros intrepidus* [11,16,85,86]. However, several other morphological differences of tyrannosauroids allow us to exclude MMNS VP-6332 from the group. For example, an extreme medial expansion of the proximal end bearing a deeply notched proximal articular surface for the third metatarsal, a dorsally bending and ventrally convex shaft in lateral view, the absence of distinct ridges on the posterior surface, and the slightly curved prominent ridge along the anterolateral corner of the distal end are features characteristic of the small to medium-bodied tyrannosauroids, such as *A. olseni*, *A. montgomeriensis*, and *M. intrepidus* [11,86] and the Delaware tyrannosauroid [16], that are not observed on MMNS VP-6332. Furthermore, the elongate, slender MMNS VP-6332 metatarsal differs from those of the robust metatarsals of the large-bodied tyrannosaurids, such as *Albertosaurus sarcophagus* (e.g., AMNH 5432), *T. rex* (e.g., AMNH 973 and FMNH PR 2081), and *G. libratus* (CMN 2120) (S3 Fig). In addition, MMNS VP-6332 can also be differentiated from tyrannosaurs by possessing a straight metatarsal with a suboval cross-section of the shaft and lacking a buttressing flange at the posterolateral border, where the second metatarsal contacts with the fourth metatarsal distally [63,87,88].

#### The third metatarsal (MT-III)

The distal half of a right third metatarsal (MSC 13139) is only preserved (Fig 5; Table 1), but the preserved portion is damaged at the mid part of the shaft (Fig 5A). The shaft is mediolaterally broad anteriorly with a flat surface that transitions proximally to become extremely thin and narrow, forming a splint bone with a slightly convex anterior surface. The cross-section of MSC 13139 is wedge-shaped and transversely wider than deep.

**Fig 5.**
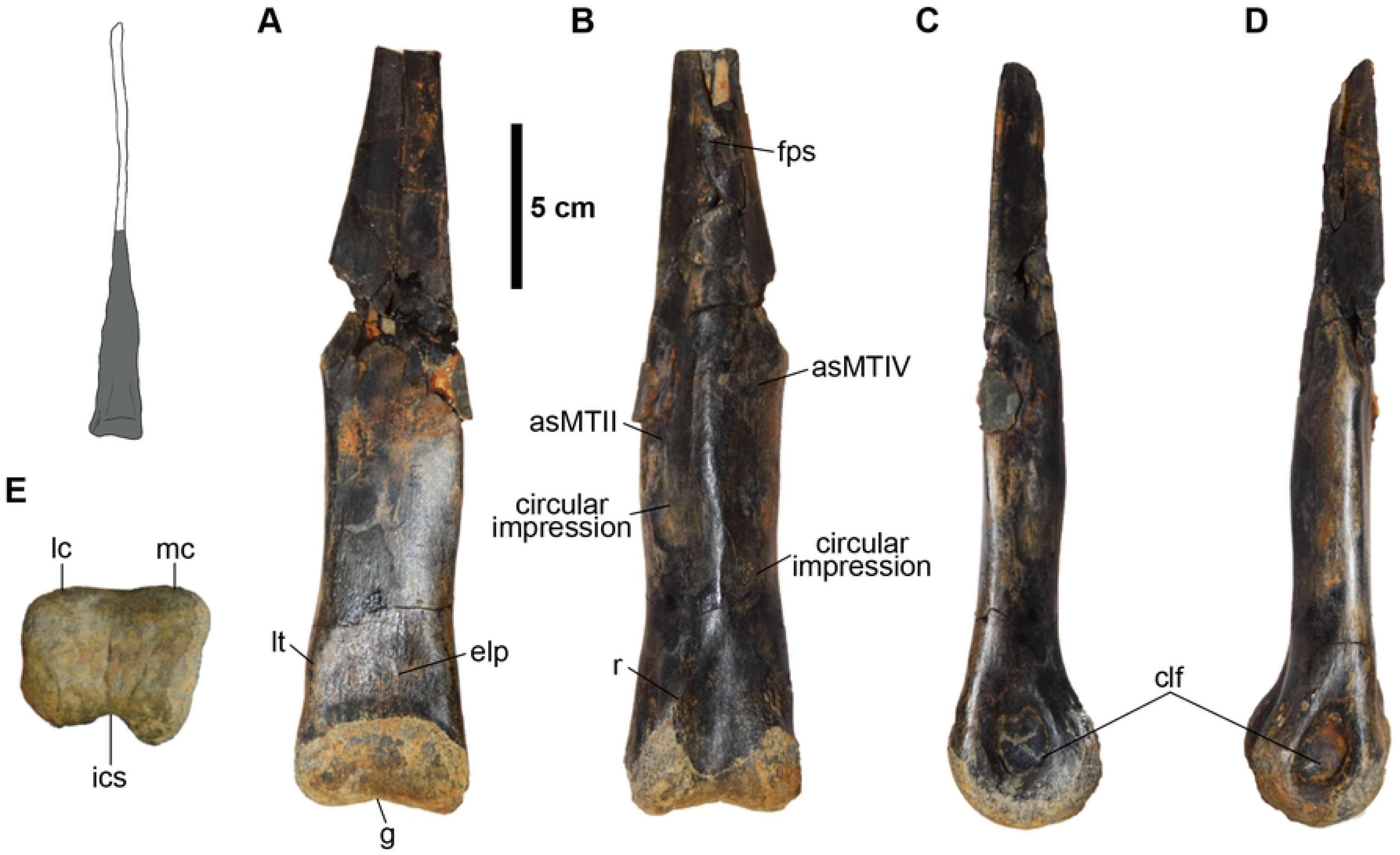
Distal half of the right third metatarsal (MSC 13139). **(A)**, anterior; **(B)**, posterior; **(C)**, medial; **(D)**, lateral; **(E)**, distal views. Interpretive illustration of *A. tugrikinensis* (MPC-D 100/130) shows the approximate location of the preserved portion of the metatarsal. Abbreviations: asMTIII, articular surface for the third metatarsal; asMTIV, articular surface for the fourth metatarsal; clf, collateral ligament fossa; elp, extensor ligament pit; fps, flat posterior surface; ics, intercondylar sulcus; lc, lateral condyle, lt, “lateral tab”; mc, medial condyle.

**Fig 6.**
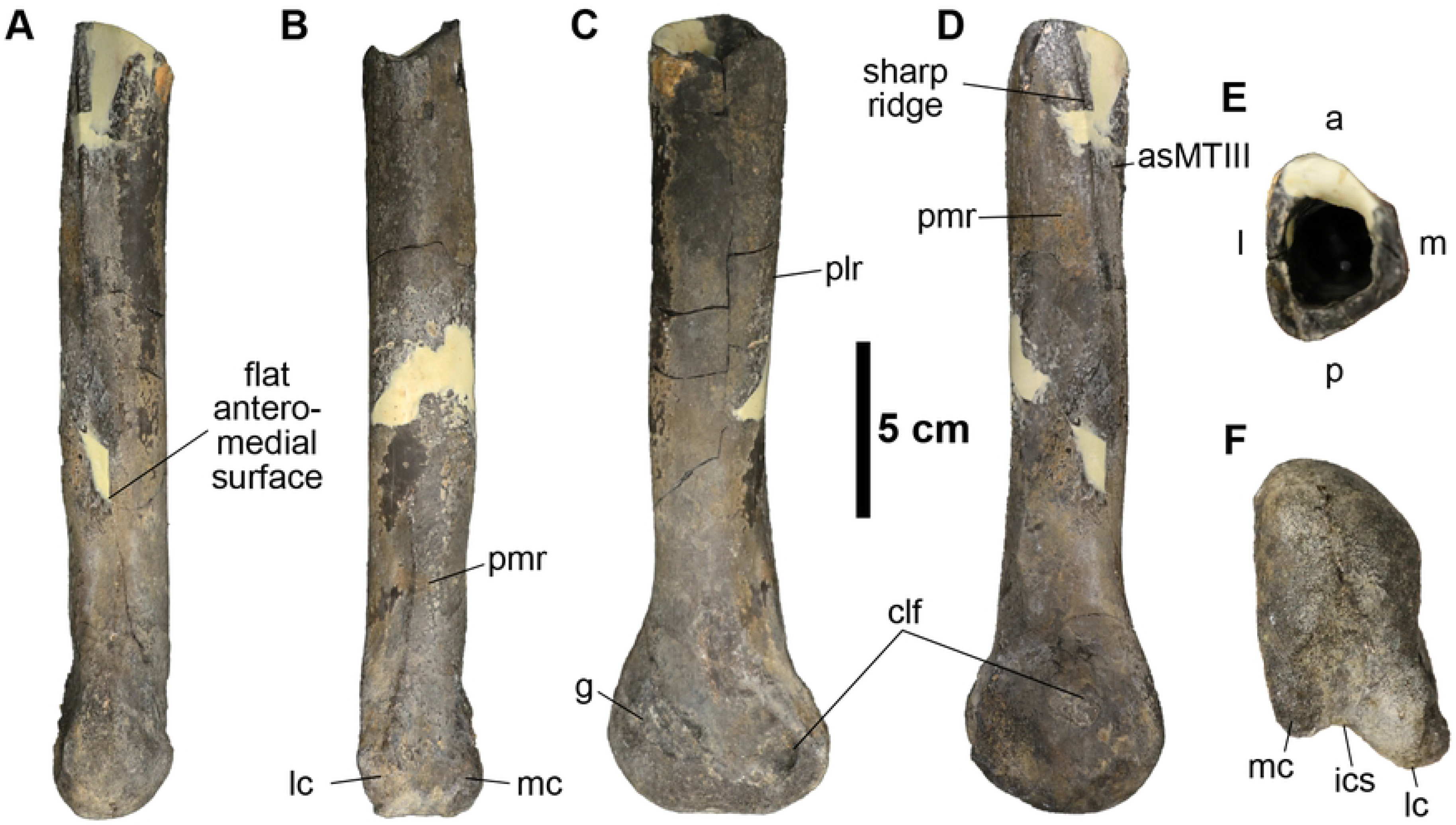
Distal half of the left fourth metatarsal (MMNS VP-6183). **(A)**, anterior; **(B)**, posterior; **(C)**, lateral; **(D)**, medial; **(E)**, proximal; **(F)**, distal views. Abbreviations: a, anterior; asMTIII, articular surface for the third metatarsal; clf, collateral ligament fossa; g, groove; ics, intercondylar sulcus; l, lateral; lc, lateral condyle; m, medial; mc, medial condyle; p, posterior; plr, posterolateral ridge; pmr, posteromedial ridge.

**Table 1.**
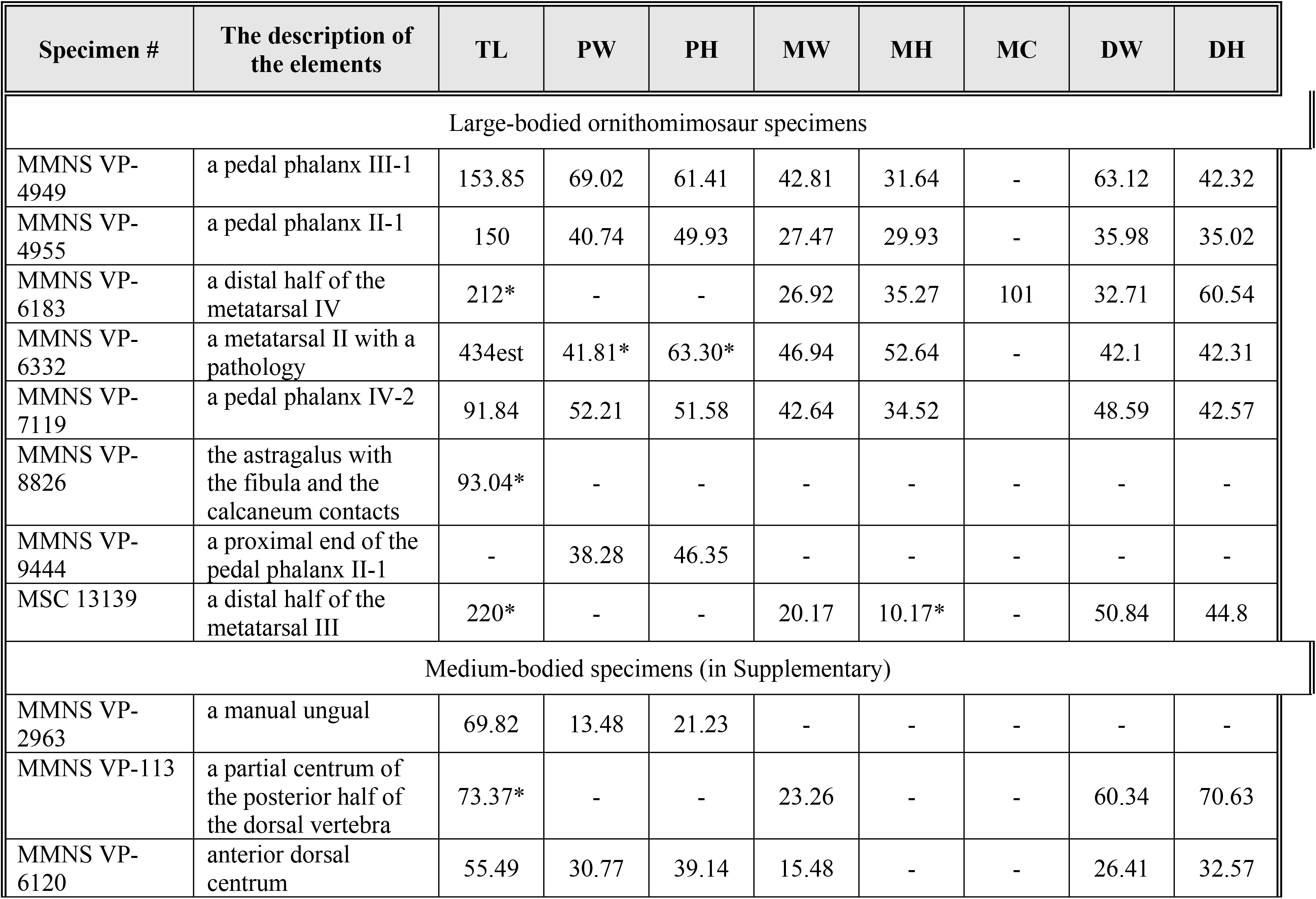
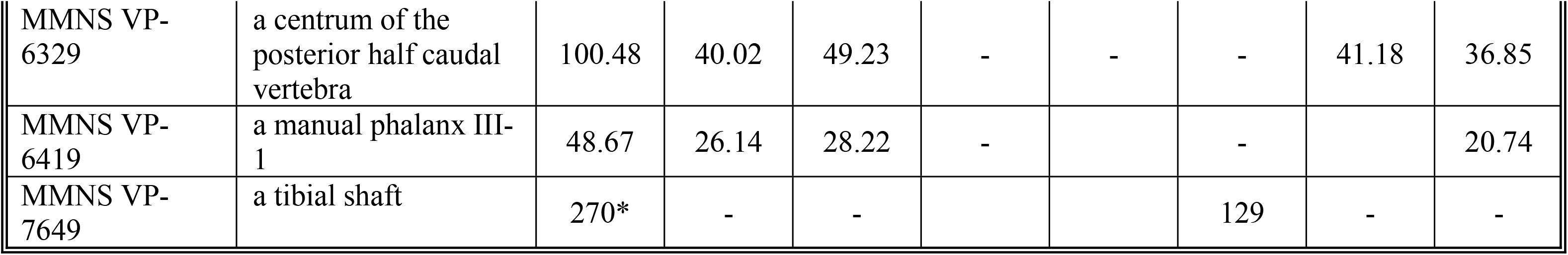
Measurements of the preserved elements of the Eutaw ornithomimosaurs. Abbreviations: (DH), distal height; (DW), distal width; (MC), midshaft circumference (least point); (MH) midshaft height (MW), midshaft width (least point); (PH), proximal height; (PW), proximal width; (TL), total length. Note that a single asterisk (*) indicate measurements are taken from the incomplete elements.

In anterior view, the medial border of the proximal-most preserved portion of the shaft is straighter when compared to the lateral one, which trends proximomedially, constricting the shaft at midlength for reception of the fourth metatarsal (Fig 5A). The latter feature is common in both arctometatarsalian and non-arctometatarsalian ornithomimosaurs (e.g., *R. evadens* and *Ornithomimus edmontonicus* [CMN 8632, ROM 797, ROM 851, and TMP 1995.110.1]). In MSC 13139, the lateral and medial articular surfaces for the adjacent metatarsals are flat and face posterolaterally and posteromedially, which is clearly indicating an arctometatarsalian foot condition as in late-diverging ornithomimosaurs (Fig 5B). Distinct, subequal circular impressions are located on each side of the posterolaterally and posteromedially facing surfaces proximal to the distal caput (Fig 5B). These depressions are presumably the attachment scars for the other metatarsals (Fig 5B). The lateral scar is positioned more distally than the medial scar, and so it is presumed that the fourth metatarsal is slightly longer than the second metatarsal, consistent with the condition observed in late-diverging ornithomimosaurs [25]. In anterior view, the lateral margin of MSC 13139 is more laterally extensive than the medial margins, which would have only slightly overhung the second metatarsal. Both lateral and medial anterior margins proximal to the distal articular caput are smooth and round, and the medial margin is slightly pinched compared to the lateral margin of *O. velox* in anterior view. In overall appearance, the morphology of the MSC 13139 shaft is similar to those of the type specimen of *O. velox* (YPM 548) and the large Gansu ornithomimid [61,65] (S4A and S4E Fig); however, it differs from the strongly concave and sinuous medial margin of the third metatarsal of *S. altus* (e.g., AMNH 5339, AMNH 5383).

Anteriorly, the outline of the lateral and medial margins of the distal end of MSC 13139 are straight and the entire distal caput is rectangular-shaped as in late-diverging ornithomimosaurs like *A. tugrikinensis, D. brevitertius, O. velox,* and *R. evadens* (S4A-D Fig), but unlike the condition in *A. fridayi*, *B. grandis*, *Q. henanensis* and the large Gansu ornithomimid, which have a more mediolaterally widened articular caputs relative to the transverse width of distal shaft (S4E-H Fig). Furthermore, MSC 13139 exhibits a small lateral process on the anterolateral surface of the distal end just proximal to the caput (Fig 5A) (“lateral tab,” sensu Zanno et al. 2011). A lateral tab is also documented on *O. velox*, *R. evadens* [74], and *A. tugrikinensis*, but it is much weaker in these taxa than the well-developed lateral tab of troodontid metatarsals (e.g., *T. sampsoni*, *Troodon formosus*, and *S. inequalis* [43,75,76]) (S4D-E Fig). A shallow, semicircular-shaped extensor ligament pit is present immediately proximal to the distal articular caput as in ornithomimosaurs (Fig 5A), which is differentiated from those of oviraptorosaurs, such as *Anzu*, *G. erlianensis* and *Heyuannia huangi* [66,67]. The degree of development is similar to late-diverging ornithomimosaurs, such as *O. velox* and *S. altus*, and unlike those of early-diverging ornithomimosaurs, such as *A. fridayi, B. grandis* and *D. mirificus*, which bears proximodistally extended deep and narrow extensor ligament pit.

The distal articular surface of MSC 13139 is smooth and non-ginglymoid with unevenly developed lateral and medial condyles (Fig 5A). The lateral and the medial distal condyles are mediolaterally subequal, but the medial condyle is posteriorly more extended and anteroposteriorly slightly taller than the lateral condyle in distal view (Fig 5E). Although this feature is present in most ornithomimosaurs, *S. altus* (CMN 930) and *O. edmontonicus* (CMN 8632) exhibit relatively equal distal condyles [91]. Moreover, the distal articular surface is mediolaterally wider than it is anteroposteriorly and bears a shallow vertical groove along the midlength (Fig 5E). The mediolaterally wider distal articular surface with anteroposteriorly more pronounced medial condyle of MSC 13139 is similar to the tyrannosauroid *A. montgomeriensis* and the caenagnathid *Elmisaurus rarus*, but it differs from the troodontid *T. sampsoni* and *S. inequalis* (Fig. S4). In anterior view, the median groove on the distal articular surface of MSC 13139 is more pronounced than other ornithomimosaurs, such as *R. evadens* and the large Gansu ornithomimid in anterior view [61,74] (S4C and S4E Fig).

In posterior view, the posterior surface of the shaft has a relatively broad flat surface (less than 1 cm wide) proximally and a posteriorly pointed surface distally (Fig 5B). The feature is similar to some ornithomimosaurs, such as *O. velox*, *G. bullatus* and the large Gansu ornithomimid [61,64,65], but it is inconsistent with *A. tugrikinensis*, which displays a sharp ridge posteriorly along the proximal shaft [62]. Distally, MSC 13139 bears the sub-equally developed lateral and medial condyles on the posterior surface of the distal caput (Fig 5B). The lateral condyle has a smooth surface, whereas a prominent longitudinal ridge is developed just proximal to the medial condyle, which extends ∼1 centimeter proximally (Fig 5B). Posteriorly, lateral and medial condyles are well-separated by a deep intercondylar sulcus like some ornithomimosaurs (e.g., *O. edmontonicus* [CMN 8632], *G. bullatus* and *Tototlmimus packardensis* [50,78]) (Fig 5E). The intercondylar sulcus is variably developed within ornithomimosaur taxa by the degree of depth [74]. For example, some ornithomimosaur taxa, such as *R. evadens* [74], display a straight outline with no intercondylar sulcus. The depth of the intercondylar sulcus on MSC 13139 is similar to *G. bullatus, Garudimimus brevipes,* the large Gansu ornithomimid, and some undescribed Asian ornithomimid specimens (pers. obs.). The depth of these sulci differs from the condition in some ornithomimosaur taxa, such as *A. fridayi*, *H. okladnikovi*, *O. edmontonicus* (CMN 8632), *O. velox* and *S. altus* (CMN 930), which bear shallow intercondylar sulci [14,61,74,80]. The lateral and medial collateral ligament fossae are equally well-developed both in size and depth in MSC 13139, similar to other theropods (Fig 5C and D). The outline of these pits is subcircular, and they are positioned at the center of each side of the distal caput. Overall, the distal end of MSC 13139 is typical of ornithomimosaurs (e.g., *A. tugrikinensis, G. bullatus, G. brevipes, O. edmontonicus, O. velox,* and *S. altus* [25,47,48]) (Fig 5A and E) albeit distinguishable from those of troodontids, another arctometatarsalian theropod clade, in which the distal articular surface is taller than wide in distal view (S5D-E Fig) [93].

MSC 13139 cannot be referred to large theropods, such as allosauroids and tyrannosaurids, nor to other medium-sized theropods, such as dromaeosaurids, troodontids, and caenagnathids, based on a number of characteristics. A proximally slender shaft with a wedge-shaped, triangular cross-section of the “arctometatarsalian condition,” shallow extensor ligament pit, and mediolaterally distally unexpanded distal articular caput in MSC 13139 clearly distinguish it from allosauroids (e.g., *Acrocanthosaurus atokensis* [NCSM 14345], and *A. fragilis*), which have a mediolaterally unconstricted transversely wide shaft and expanded distal articular caput with a deep extensor ligament pit [94,95]. These features are also present in tyrannosaurids (e.g., *A. sarcophagus*, *Daspletosaurus torosus,* and *T. rex* [82]). However, several other morphological features of tyrannosaurids, such as a strongly deviated (medially bulging) medial anterior margin, anteroposteriorly taller than mediolaterally wider shaft at midshaft, deep external ligament pit, and subrectangular distal articular surface without an intercondylar sulcus, differentiate the MSC 13139 metatarsal from tyrannosaurids. Most of these characteristics are present even in early ontogenetic stages in tyrannosaurids, based on our observations of a juvenile specimen of *T. bataar* (MPC-D 107/7), thus they are reliable diagnostic features. Similar to the slender metatarsals of more early-diverging, non-tyrannosaurid tyrannosauroids, such as *A. olseni, A. montgomeriensis*, and *G. libratus*, are the anteriorly unexpanded, straight distal articular caput in lateral view, a wedge-shaped cross-section of the shaft with flat articular surfaces for the adjacent metatarsals, a larger medial condyle, and centrally positioned, equally developed collateral ligament fossae [11,85,97]. However, the presence of a deep extensor ligament pit, a distinct demarcating lip on the posterior articular surface of the distal condyle, and a single, ungrooved distal articular surface with a lack of the intercondylar sulcus of these same taxa exclude MSC 13139 from tyrannosauroids.

MSC 13139 also cannot be referred to dromaeosaurids, oviraptorosaurs, and troodontids. MSC 13139 is differentiated from dromaeosaurids and some oviraptorosaurs by an arctometatarsalian condition with a non-ginglymoid articular surface [81,82,84,98]. On the other hand, resemblance to troodontids is seen in an arctometatarsalian third metatarsal with asymmetrical distal condyles (exhibiting a larger medial condyle), straight lateral and convex medial borders in distal view, a shallow sulcus on the articular surface, a shallow semi-circular extensor ligament fossa, and a transversely slight narrowing shaft directly proximal to the distal caput on the anterior surface of MSC 13139 (e.g., *G. mongoliensis*, *Stenonychosaurus inequalis, T. sampsoni* and *Zanabazar junior*) [57,58,89,99]. However, MSC 13139 can be differentiated from troodontids by a distal end that is mediolaterally wider than anteroposteriorly tall in distal view, a less pronounced lateral tab on the anterolateral margin of the distal caput, and a weakly developed intercondylar sulcus on the posterior surface. A subequally proportioned distal articular surface in distal view with an anteroposteriorly more extended medial condyle, and a posteriorly demarcated distinct lip, are troodontids features (e.g., *S. inequalis* and *T. sampsoni*) that can be used to rule out the affinity of that group with MSC 13139 [57,90] (S5D-E Fig).

#### The fourth metatarsal (MTIV)

The distal half of a left fourth metatarsal (MMNS VP-6183) bears a well-preserved distal articular surface, mediolaterally compressed distal caput, laterally flared distal condyle, and articular surface for the third metatarsal along the medial surface of the shaft (Fig 6). The shaft is straight and slender relative to its width, which differs from the robust tyrannosaurids’ metatarsals (e.g., *A. sarcophagus, G. libratus,* and *T. rex*) [63,87,97], but not from slender-footed tyrannosauroid taxa (e.g., *A. olseni,* and *M. intrepidus*) [85,86]. The preserved distal shaft is anteroposteriorly taller than the mediolateral width (Fig 6, Table 1). As in most late-diverging ornithomimosaurs, the shaft exhibits flat anteromedial (articular surface for the third metatarsal) and posteromedial surfaces, separated by a sharp ridge, and a slightly convex lateral surface, displaying a subtriangular shape in proximal cross-sectional view (Fig 6E). This contrasts with the condition of early-diverging ornithomimosaurs, such as *A. fridayi* and *G. brevipes*, which display a subrectangular shaft in cross-section [14,100]. In lateral view, the shaft slightly narrows just proximal to the distal articular caput, creating a slight anterior arch (Fig 6C). A similar arching is observed in tyrannosauroids such as *A. sarcophagus* (ROM 807)*, A. montgomeriensis,* and *M. intrepidus* [11,86]; however, it is much more pronounced in those taxa than in MMNS VP-6183. The posterior surface of the corresponding region of MMNS VP-6183 is rugose and bears two distinct ridges--posteromedial and posterolateral (Fig 6C-D). The posteromedial ridge originates proximal to the lateral condyle and extends to the posteromedial edge, confluent with the posteromedial margin of the shaft (Fig 6B). The posterolateral ridge is slightly stronger and broader than the posteromedial and gradually fades just proximal to the distal articular caput. This ridge is much weaker than the condition observed in slender-footed tyrannosauroids (e.g., *M. intrepidus* [72]).

The distal articular caput of MMNS VP-6183 is slightly rotated counter-clockwise in distal view (Fig 6F). The distal articular surface is taller than wide due to the extreme anteroposterior expansion of the distal caput, visible in medial or lateral views (Fig 6C-D). Although anterior expansion is more commonly seen in ornithomimosaurs than in tyrannosauroids, the degree of the development is variable among early-diverging species of ornithomimosaurs, such as *A. fridayi* and *G. brevipes*, bearing an anteriorly expanded distal caput [14,100], as well as late-diverging taxa bearing an anterior margin more in line with the anterior shaft in lateral view (e.g., *O. velox, S. altus* (AMNH 5339) and *R. evadens* [47,60,87]). The smooth, non-ginglymoid, anteroposteriorly taller than wide distal articular surface of MMNS VP-6183 resembles that of the tyrannosauroid *M. intrepidus*; however, the distal caput is not anteriorly expanded in *M. intrepidus*, nor in other tyrannosaurs (e.g., *A. sarcophagus* and *A. montgomeriensis*), which are characterized by a straight anterior margin over the distal caput in lateral view [11,86]. Although the tyrannosauroid *D. aquilunguis* exhibits a similar condition to MMNS VP-6183, the slenderness of MMNS VP-6183, with its anteroposteriorly taller than wide shaft, is distinct from *D. aquilunguis* [102].

The medial collateral ligament fossa of MMNS VP-6183 is circular and larger than the proximodistally elongated lateral fossa (Fig 6C-D). Whereas the medial collateral fossa is centered on the distal caput, the lateral collateral ligament fossa is located close to the posterior margin of the lateral condyle. MMNS VP-6183 bears a prominent groove on the anterolateral surface of the distal caput (Fig 6C). Although this groove is similar to that reported on the corresponding surface of the fourth metatarsal of the tyrannosauroid *M. intrepidus* ([72], fig. 3g-h), it is much shallower and does not extend to the collateral ligament fossa, which is distinguishable from the condition in *M. intrepidus*. Posteriorly, the distal caput has two unevenly developed distal condyles; the lateral condyle is larger than the medial condyle, as in *M. intrepidus*, yet unlike the condition of other tyrannosauroids (e.g., *A. montgomeriensis*, *G. libratus,* and *T. rex* [11,49]) (Fig 6B and F). The lateral and the medial condyles are separated by a weakly developed intercondylar sulcus in distal view, similar to that observed on the Delaware tyrannosauroid metatarsal [16], but in contrast to the condition in *A. montgomeriensis* ([11], fig. 19g). The proximodistally straight shaft and absence of a posteriorly backswept distal caput allows us to rule out a tyrannosauroid affinity for MMNS VP-6183 (e.g., [58,72]).

#### Pedal phalanges

Four pedal phalanges are preserved in the Eutaw ornithomimosaur assemblage, but appear to belong to different individuals (Fig 7). Two of them are associated with left and right sides of the first phalanx (MMNS VP-9444 and MMNS VP-4955), one is a first phalanx of third digit (MMNS VP-4949), and one is a second phalanx of fourth digit (MMNS VP-7119).

**Fig 7.**
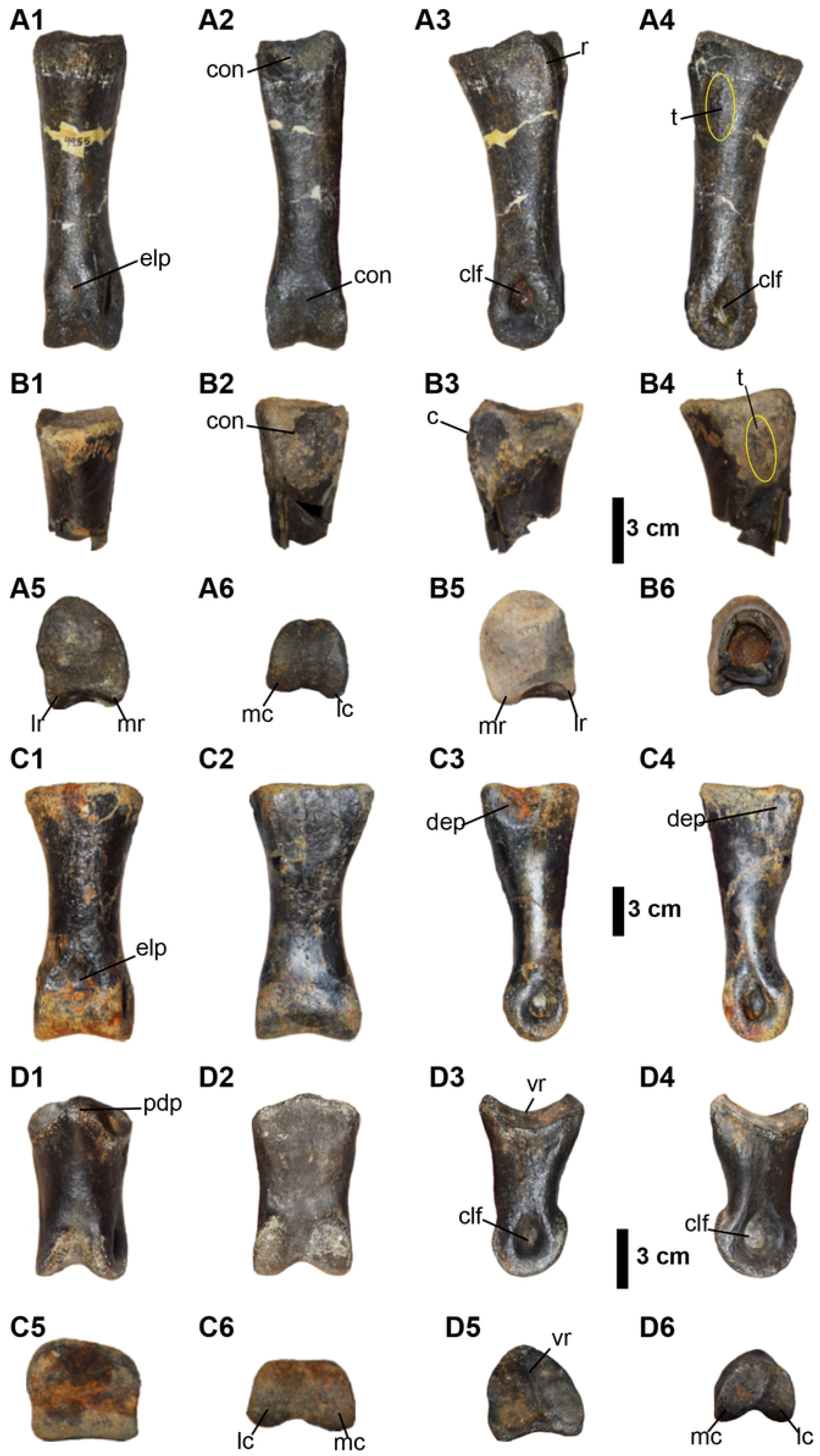
Pedal phalanges of Eutaw ornithomimosaurs. (A1-A6) the first phalanx of the second digit of left foot, (MMNS VP-4955); (B1-B6) the first phalanx of the second digit of right foot, (MMNS VP-9444); (C1-C6) the first phalanx of the third digit of right foot, (MMNS VP-4949); (D1-D6) the second phalanx of the fourth digit of left foot, (MMNS VP-7119). (A1-D1) anterior, (A2-D2) posterior, (A3-D3) lateral, (A4-D4) medial, (A5-D5) proximal, and (A6-D6) distal views. Abbreviations: c, curved ridge; clf, collateral ligament fossa; con, concavity; dep, depression on the lateral and medial surfaces of the proximal end; elp, extensor ligament pit; lr, lateral ridge; mr, medial ridge; pdp, posterodorsal process; r, ridge; t, tubercle; vr, vertical ridge.

#### Pedal phalanx (PII-1)

Two phalanges, one partial (MMNS VP-9444) and one complete (MMNS VP-4955), are referable to the first phalanx (PII-1) of the second pedal digit (Fig 7A and B). This referral is based on the length/width ratio of the phalanx, greater height than width, singular concave articular surface, and a well-developed intercondylar groove on the posterior surface of the proximal end. MMNS VP-9444 (a right phalanx) and MMNS VP-4955 (a left phalanx) are subequal in size and derived from individuals of similar size or possibly from a single individual. The following description uses both specimens in combination. In general proportions, MMNS VP-4955 is proximodistally three times longer than the height of its proximal articular surface, as in ornithomimosaurs (Table 1), and unlike PII-1 of tyrannosaurids and oviraptorosaurs, which are proximodistally shorter relative to the height of the proximal end. In dorsal view, the shaft of MMNS VP-4955 is asymmetrical with somewhat concave lateral and straight medial margins (Fig 7A1) closely resembling the condition observed in most ornithomimosaurs. This is in contrast to other large theropods, such as carcharodontosaurids, caenagnathids and tyrannosaurids [63,87,94,95,103], and decidedly distinct from the pedal PII-1 of dromaeosaurids (e.g., *D. albertensis* (AMNH 5356) [90]), and therizinosauroids (e.g., *F. utahensis* [45]), which possess a subequally and mediolaterally constricted shaft in anterior view (Fig 7A1 and A2).

The proximal end of MMNS VP-4955 is taller than wide, exhibiting a single concave articular surface in proximal view (Fig 7A5, Table 1). In proximal view, the margin of the proximal articular surface is anteromedially round, laterally straight, and slightly concave posteriorly, which is somewhat distinguishable from other theropods (Fig 7A5 and 7B5). Although the anteriorly round border of the proximal articular surface of MMNS VP-4955 is similar to tyrannosaurids (e.g., *T. rex* [FMNH PR 2081], *A. sarcophagus* [CMN 11315], and *G. libratus* [CMN 2120]) [63,87,96], the straight lateral and round medial borders of the proximal articular surface is different from the condition in tyrannosaurids, which display laterally round and medially angled borders in proximal view. The outline of the proximal articular surface of MMNS VP-4955 is somewhat similar to that of the large theropod *Allosaurus fragilis* [94]; however, it is easily differentiated by the proportions of its relative proximodistal length. Both lateral and medial articular borders of the proximal end are straight in lateral and medial views (Fig 7A1 and 7A3-A4), as in ornithomimosaurs. This is unlike dromaeosaurids, such as *D. antirrhopus* (YPM 5205), *D. albertensis* (AMNH 5356), and *V. mongoliensis*, which exhibit highly concave lateral and medial articular borders and a median vertical ridge that is visible in lateral and medial views. On the posterior surface, there are two distinct ridges along the lateral and medial borders of the proximal heel (Fig 7A5). These ridges are separated by a deep sulcus as in other theropods, such as ornithomimosaurs and tyrannosaurids [14,25,62,74,75,96], but unlike those of oviraptorosaurs and therizinosauroids, which typically exhibit a relatively flat surface at this region. The medial ridge is mediolaterally wide and stouter than the lateral ridge (Fig 7A5 and 7B5). Likewise, in the holotype specimen of *O. velox* [61], both lateral and medial ridges on the heel are curved in lateral and medial views (Fig 7A3-A4), unlike the condition in Asian taxa, such as *Aepyornithomimus tugrikinensis* and *G. brevipes*. *A. tugrikinensis* differs by exhibiting a straight lateral ridge (fig. 4c and 4d, [48]); *G. brevipes* is distinguished by a lateral ridge on the heel that is squared in lateral view (fig. 17c, [86]). There is a weak, proximodistally extended tubercle present on each lateral and medial surfaces of the proximal end of MMNS VP-4955 (Fig 7A4). A similar tubercle is also present on the corresponding surfaces of MMNS VP-9444, suggesting that this tubercle is not a pathology, and the morphologically similar tubercle is also found on the holotype specimen of *O. velox*. This contrasts with *A. tugrikinensis*, which exhibits a more laterally tilted posterolateral border of the proximal end in anterior view.

Furthermore, unlike the relatively straight anterior surface of tyrannosaurids, the anterior surface of MMNS VP-4955 gently slopes from the proximal end to the distal end in lateral view (Fig 7A3) until it reaches the distal caput as in ornithomimosaurs. Overall, the outline of the proximal articular surface of MMNS VP-4955 is consistent with most ornithomimosaurs, such as *A. tugrikinensis*, *G. brevipes*, *O. velox*, and the Bissekty taxon [61,62,100,105]. However, it is different from PII-1 of the early-diverging ornithomimosaur *A. fridayi*, which exhibits a mediolaterally wide and nearly round proximal articular surface in proximal view [14]. The cross-section of the shaft is round as in other ornithomimosaurs (Fig 7B6).

On the anterior surface, just proximal to the distal condyles, MMNS VP-4955 bears a shallow, weakly developed external ligament pit (Fig 7A1), which also resembles those of late-diverging ornithomimosaurs, such as *A. tugrikinensis, G. bullatus, O. edmontonicus* (ROM 851), *O. velox, Sinornithomimus dongi*, and *S. altus* (CMN 930 and TMP 90.26.01) [24,25,61,62,64,106], but is unlike those of early-diverging ornithomimosaurs *A. fridayi* and *Nqwebasaurus thwazi* that exhibit a relatively deep external ligament pit on the corresponding surface of the phalanges [14,107]. Posteriorly, the MMNS VP-4955 shaft is relatively flat, the only concavity present is slight and lies between the lateral and medial ridges proximally and between the condyles distally, as in other ornithomimosaurs (Fig 7A2 and 7A5-A6). This is unlike the condition in oviraptorosaurs (e.g., *G. erlianensis* and *Oksoko avarsan* [53,94]).

The distal end of MMNS VP-4955 exhibits subequally developed lateral and medial condyles, divided by a shallow midline groove distally (Fig 7A1). The lateral condyle is slightly constricted mediolaterally and is anteroposteriorly longer than the medial condyle in distal view (Fig 7A6). The anteroposterior height and mediolateral width of the distal articular surface are subequal, which is differentiated from transversely wider than tall distal articular surfaces of tyrannosaurid taxa, such as *T. rex* (FMNH PR 2081) and *G. libratus* (FMNH PR 2211). In this respect, MMNS VP-4955 closely resembles those of ornithomimosaurs (Fig 7A6, Table 1). In addition, the borders of the lateral and medial distal condyles are closely positioned anteriorly, exhibiting a mediolaterally constricted articular surface in distal view. The feature is different from allosaurids, caenagnathids (e.g., *G. erlianensis*), and tyrannosaurids, as well as some medium-sized theropods, such as dromaeosaurids (e.g., *D. antirrhopus* and *D. albertensis*) and oviraptorosaurs (e.g., *O. avarsan*), which exhibit more widely placed distal condyles with a highly ginglymoid articulation [81,104,108]. Moreover, the lateral and medial condyles of MMNS VP-4955 are relatively flat and are not expanded anteriorly as they are in dromaeosaurids like *Achillobator giganticus*, *A. mongoliensis*, and *D. antirrhopus,* and troodontids like *S. inequalis* (CMN 1650) and *T. sampsoni* in lateral view [57,82,98,104]. The collateral ligament fossae of MMNS VP-4955 are well developed; the lateral collateral ligament fossa is larger and deeper than the medial one (Fig 7A3-A4). The condition is similar to ornithomimosaurs, but it is unlike tyrannosaurids (e.g., *A. sarcophagus* [CMN 11315], *G. libratus* [FMNH PR 2211], *T. rex* [FMNH PR 2081 and BMRP 2002.4.1]), which exhibit subequal-sized collateral ligament fossae. Although the morphology of the distal articular surface of MMNS VP-4955 is more consistent with ornithomimosaurs than other theropods, it also differs from specific ornithomimosaurs. For example, whereas MMNS VP-4955 bears equally pronounced distal condyles in anterior view, *A. tugrikinensis* has a more distally extended medial condyle (Chinzorig et al. 2017).

#### Pedal phalanx (PIII-1)

MMNS VP-4949 is referable to the first phalanx (PIII-1) of the third digit based on its relative symmetry, wider than tall proportions, single and slightly concave proximal articular surface, and weakly or non-grooved, non-ginglymoid distal articular surface with subequally developed condyles (Fig 7C). Both proximal and distal articular surfaces are slightly eroded; however, the phalanx is essentially complete except for a small portion of the anterolateral surface of the distal half (Fig 7C1). MMNS VP-4949 is proximodistally slender and elongate relative to the mediolateral width of the proximal end in anterior view (Fig 7C1). Comparatively, it is approximately two or three times longer than those of medium-sized ornithomimosaurs (Table 2) and one to two times longer than those of large-bodied ornithomimosaur taxa (e.g., 171% the length of PIII-1 in *G. bullatus* [MPC-D 100/11] and 169% of *A. fridayi*) and approaches the size of *D. mirificus* (Table 2). A proximodistally slender PIII-1 is similar to those of the noasaurine *Vespersaurus paranaensis* [109], *D. antirrhopus* [81], and the tyrannosauroids *A. olseni* and *Gorgosaurus*. The length/width ratio of the proximal articular surface of MMNS VP-4949 is subequal, forming a subrectangular outline in proximal view, compared to the distal articular surface, which is much wider than its anteroposterior length (Fig 7C5). The proximal articular surface is anteriorly convex, posteriorly straight, and exhibits slightly notched lateral and medial borders in proximal view (Fig 7C5). MMNS VP-4949 also has a relatively straight articular border in anterior view (Fig 7C1), which differentiates it from the anteriorly round border of the proximal articular ends of tyrannosaurids, such as *T. rex* (e.g., FMNH PR 2081), *T. bataar* (e.g., MPC-D 107/2), as well as allosaurids *A. fragilis* (YPM 1930). In this respect MMNS VP-4949 is somewhat similar to the PIII-1 of the large caenagnathids *A. wyliei* (CMN 78000) and *G. erlianensis*, as well as the therizinosauroid *F. utahensis* [59,67]. When viewed proximally, the anterior aspect of the proximal surface of MMNS VP-4949 is slightly broadened mediolaterally, compared to the posterior half (Fig. 7C5). This feature may be unique because it is not found on ornithomimosaurs, tyrannosaurids (e.g., *T. rex* [FMNH PR 2081] and *G. libratus* [FMNH PR 2211]), allosaurids (e.g., *A. fragilis* [YPM 1930]), noasaurids (e.g., *V. paranaensis*), or oviraptorosaurs. However, several features differentiate MMNS VP-4949 from many of these theropods, including the presence of a nearly straight margin of the proximal articular end in lateral view and a mediolaterally wide distal articular end. In addition, MMNS VP-4949 lacks an extremely deep external ligament pit and ginglymoid distal articulation (Fig 7C1 and S6 Fig), which is characteristic of many of the aforementioned incidents.

**Table 2.** Length comparisons of different pedal phalanges between the Eutaw ornithomimosaurs and other described ornithomimosaurs. Note that a single asterisk (*) indicates the estimated value and dash (-) indicates unknown elements.

| Species names | Specimen # | Pedal phalanx (II-1), in mm | Pedal phalanx (III-1), in mm | Pedal phalanx (IV-2), in mm |
| --- | --- | --- | --- | --- |
| *Aepyornithomimus tugrikinensis* | MPC-D 100/130 | 59.08 | 52 | 24 |
| *Anserimimus planinychus* | MPC-D 100/300 | 65 | 56 | 17 |
| *Archaeornithomimus asiaticus* | AMNH 6565 | 71 | - | - |
| *Arkansaurus fridayi* | 74-16-3-5 | 102 | 91 | 71 |
| *Beishanlong grandis* | FRDC-GJ (06) 01-18 | 120 | - | - |
| *Deinocheirus mirificus* | MPC-D 100/127 | 192 | 165 | 89.15 |
| *Dromiceiomimus brevitertius* | CMN 12068 | 67.4 | 56 | 21 |
| *Dromiceiomimus brevitertius* | CMN 12069 | 57.3 | 27.8 | 20.2 |
| *Dromiceiomimus brevitertius* | ROM 797 | 53 | - | 33 |
| *Dromiceiomimus brevitertius* | ROM 852 | 80 | - | 27 |
| *Gallimimus bullatus* | MPC-D 100/10 | 32 | 31 | 13 |
| *Gallimimus bullatus* | MPC-D 100/11 | 102 | 90 | 43 |
| *Gallimimus bullatus* | MPC-D 100/52 | 59 | 57 | 11 |
| *Gallimimus bullatus* | ZPal MgD-I/8 | - | 97 | 50 |
| *Gallimimus bullatus* | ZPal MgD-I/94 | 45 | 44 | 19 |
| *Garudimimus* | MPC-D 100/13 | 63 | 59 | 35 |
| *Harpymimus okladnikovi* | MPC-D 100/29 | 72 | 67 | 34 |
| *Nqwebasaurus thwazi* | AM 6040 | 20 | - | - |
| *Ornithomimus edmontonicus* | CMN 8632 | 78 | - | 23.5 |
| *Ornithomimus edmontonicus* | ROM 851 | 78.8 | 72.3 | 35 |
| *Ornithomimus edmontonicus* | TMP 95.110.1 | 68.5 | - | 22 |
| *Paraxenisaurus normali* | BENC 2/2-001 | 115 | - | - |
| *Sinornithomimus dongi* | IVPP V11797-10 | - | 50.3 | 21.8 |
| *Struthiomimus altus* | AMNH 5375 | 82 | 76 | 28 |
| *Struthiomimus altus* | AMNH 5257 | 83 | 87 | 31 |
| *Struthiomimus altus* | AMNH 5339 | 85 | - | 26 |
| *Struthiomimus altus* | CMN 930 | 75 | 67 | 21 |
| *Struthiomimus altus* | TMP 90.26.1 | 94.2 | 80 | 26.9 |
| *Struthiomimus altus* | UCMZ (VP) 1980.1 | 92 | 81 | 27 |
| *Rativitas evadens* | ROM 1790 | 63.9 | 63.3 | 19.6 |
| *Gallimimus indet.* | MPC-D 100/138 | 92 | 85.3 | 29 |
| *Ornithomimus sp.* | MNA Pl.1762A | 76 | - | 26 |
| *Ornithomimidae indet.* | MPC-D 100/121 | 65 | - | 23 |
| Large Gansu ornithomimid | IVPP V 12756 | 100.23 | - | 84 |
| **Eutaw ornithomimosaur** | **MMNS VP-4955** | 150 | 140* | 110* |
| **Eutaw ornithomimosaur** | **MMNS VP-4949** | 164* | 153.85 | 124* |
| **Eutaw ornithomimosaur** | **MMNS VP-7119** | 146* | 136* | 91.84 |

Anteriorly, the shaft of MMNS VP-4949 is straight (Fig 7C1), which is slightly constricted mediolaterally and wider than tall, forming an oval shape in cross-section. The oval-shaped cross-section of MMNS VP-4949 is like those of ornithomimosaurs, but unlike large theropods, such as *A. fragilis* (YPM 1930), *G. erlianensis*, and *T. rex* (FMNH PR 2081), which exhibit a taller than wide shaft with round or square cross-sections (S6I-L Fig).

In lateral view, the anterior surface of MMNS VP-4949 is straight, as in *T. rex* (e.g., FMNH PR 2081) and *T. bataar* (e.g., MPC-D 107/2), but the morphologies of the lateral and medial borders of the proximal ends differ from these theropods in anterior view. For example, both lateral and medial sides of the proximal end are straight in MMNS VP-4949 and other ornithomimosaurs (Fig 7C1), whereas tyrannosaurids have mediolaterally broadened, more outwardly inclined lateral and medial borders relative to the midshaft (S6 Fig). The latter feature is also seen in some caenagnathids, such as *G. erlianensis* [67].

Each side of the posterolateral and posteromedial surfaces of the proximal end of MMNS VP-4949 bear depressions (Fig 7C3-C4). Similar depressions are seen in ornithomimosaurs, such as *A. tugrikinensis, G. brevipes, H. okladnikovi* and *R. evadens* [62,74,80,100]. The lateral depression is more distinct and deeper than the medial one. The lateral depression is similarly deep in *G. brevipes*; however, in the latter taxon it forms a crescentic shape rather than a groove [100]. The presence of the distinct depressions seems to differentiate this taxon from *A. fridayi*, although the corresponding surfaces of *A. fridayi* are incomplete [14]. As in other ornithomimosaurs, the posterior surface of MMNS VP-4949 is relatively flat and straight in lateral view (Fig 7C2-C4). Although MMNS VP-4949 displays a rugose surface on the posterior aspect of the proximal end as in allosauroids, acrocanthosaurs and tyrannosauroids, it is much smoother than observed in these taxa (Fig 7C2).

Distally, MMNS VP-4949 bears a shallow extensor ligament pit on the anterior surface immediately proximal to the distal caput, which is wider distally than proximally, forming a triangular shape in anterior view (Fig 7C1). Although the feature of the proximally narrowed, shallow extensor ligament pit of MMNS VP-4949 is similar to caenagnathids, such as *Anzu* sp. (CMN 78000) and *Chirostenotes elegans* (CMN 8538), a lack of the mediolaterally subequal distal end relative to the proximal end and a non-ginglymoid articular surface differentiates MMNS VP-4949 from these taxa. Although some tyrannosauroids (e.g., *A. sarcophagus* [CMN 11315], *A. olseni* [MPC-D 100/51], and *G. libratus* [FMNH PR 2120]), also possess a proximodistally long and shallow extensor ligament pit. However, certain features allow us to rule out referral to this group, including the anteriorly expanded distal condyles relative to the proximal width and lack of a well-developed notch immediately proximal to the anterior surface of the distal caput, which are present on these same tyrannosauroids.

Distally, the lateral condyle is the smaller of the two and is slightly inclined dorsolaterally compared to the medial condyle. Moreover, the distal articular surface of MMNS VP-4949 is weakly concave and divided by a shallow vertical sagittal ridge in anterior view (Fig 7C1). The weakly concave distal articular surface is somewhat similar to that of tyrannosauroids and intermediately developed when compared to caenagnathoids. For example, the distal articular surface of the phalanx PIII-1 of *Citipes* is straight and lacks the sulcus in anterior view, whereas *Anzu* bears much greater concavity than MMNS VP-4949. However, in caenagnathoids, such as *C. elegans* (CMN 8538) and *Citipes* [79,110], the posterior margins of the lateral and medial distal condyles are not visible, whereas they are visible in MMNS VP-4949.

MMNS VP-4949 has comparatively large and equally deep collateral ligament fossae (Fig 7C3-C4), which are different in morphology and position. The lateral collateral ligament fossa is subcircular, slightly larger than the medial fossa, and is positioned close to the anterior border of the condyle in lateral view. The medial collateral ligament fossa, on the other hand, is lachrymiform, proximodistally elongated, and is positioned centrally on the condyle, bearing a distinct groove proximally (Fig 7C3-C4). This differs from the condition in tyrannosaurids, which exhibit morphologically similar and subequally-sized collateral ligament fossae on both sides of the distal caput [63,96], yet is similar to those of ornithomimosaurs, such as *G. brevipes, O. edmontonicus* (CMN 8632), and *S. altus* (CMN 930) [91].

#### Pedal phalanx (PIV-2)

MMNS VP-7119 is a complete and well-preserved phalanx (Fig 7D). Proximodistally longer than transversely wide, anteroposteriorly taller than wide, highly asymmetrical, and possessing strongly ginglymoid articular surfaces are features indicating that this specimen represents the second phalanx of left digit IV.

The articulation of the proximal end is divided into two subequal concave surfaces by a strong vertical ridge, visible in lateral view (Fig 7D3) as in *D. antirrhopus* (YPM 5205). This is in contrast to tyrannosaurids, such as *G. libratus* (FMNH PR 2211) and *T. rex* (FMNH PR 2081), which display a weakly developed ridge. As in ornithomimosaurs, the posterior process of the proximal articular surface is slightly more extended proximally than the anterior process in lateral view (Fig 7D4), which is in contrast to the tyrannosaurids condition of subequal development of processes. In anterior view, the anterior process is pointed proximally as in most ornithomimosaurs and dromaeosaurids, such as *D. antirrhopus* [111]; this differs from the rounded condition in tyrannosaurids. However, MMNS VP-7119 can be differentiated from *D. antirrhopus* by its asymmetrical and relatively straight shaft mediolaterally. The lateral border of the proximal articular end of MMNS VP-7119 is straight in proximal view, which closely resembles the condition in ornithomimosaurs but contrasts with Tyrannosauroidea (and possibly Allosauridae), which usually exhibit a sub-rectangular posteromedial border [63,87,96] (although the latter feature is less pronounced in *G. libratus* [FMNH PR 2211] than *A. sarcophagus* [CMN 11345], and *T. rex* [FMNH PR 2081]). Nonetheless, MMNS VP-7119 can be distinguished from tyrannosaurs, namely *T. rex* (FMNH PR 2081) and *G. libratus* (FMNH PR 2211), in that the proximal articular surface is slightly inclined laterally, exhibiting laterally and medially convex, and posteriorly concave borders in proximal view (Fig 7D5) [63,87].

The distal condyles are only slightly wider transversely in dorsal view compared to the shaft (Fig 7D3-D4). The condition is like that of dromaeosaurids, such as *D. antirrhopus*, but unlike those of tyrannosauroids, which have distal condyles that are greatly expanded mediolaterally relative to the shaft. The shaft of MMNS VP-7119 is robust and nearly straight mediolaterally and anteroposteriorly, as in ornithomimosaurs, and somewhat like *A. fragilis* (Fig. 7D1 and D3). In this respect, it differs from those of dromaeosaurids and tyrannosaurids, which have a shaft that strongly constricts near the distal condyles in lateral view However, *A. fragilis* can be differentiated from MMNS VP-7119 by the degree of difference between the mediolateral and anteroposterior width of the shaft, which is much greater in *A. fragilis* [94]. Laterally, MMNS VP-7119 exhibits a small, but prominent, obliquely oriented groove anteroposteriorly, on the posterolateral and posteromedial surfaces of the proximal end (Fig 7D3-D4).

Posteriorly, the proximal half of the posterior surface is slightly concave, with a rugose surface and weakly developed lateral and medial ridges. The shaft of MMNS VP-7119 is semicircular in cross-section due to a flat posterior surface.

The distal articular surface of MMNS VP-7119 is ginglymoid and bears unequally developed lateral and medial distal condyles, divided by a strong vertical ridge in distal view (Fig 7D6). Anteriorly, these condyles are closely positioned to each other in anterior view and laterally inclined in distal view as in ornithomimosaurs (Figs 7D1 and 7D6) but unlike the borders of *A. fragilis*, *G. libratus*, and *T. rex*, as well as those of medium-sized theropods, such as *D. antirrhopus*. The mediolateral width of the condyles is approximately the same; however, the medial condyle is slightly larger and taller anteroposteriorly than the lateral condyle. In addition, whereas the axis of the lateral condyle is relatively perpendicular to the transverse width, the medial distal condyle is obliquely oriented, with an anteromedial/posterolateral axis. This differs from the condition in tyrannosaurids such as *T. rex* (FMNH PR 2081), which has a straight medial condyle in distal view [63].

#### Bone microstructure

We examined osteohistological samples from two hind limb elements each belonging to a different ornithomimosaur size class occurring in the Eutaw assemblage. The bone matrix in both primarily preserve a woven-parallel complex with plexiform and reticular vascularization, and more isolated regions of longitudinal vascularization in the outer cortex. An external fundamental system (EFS) is absent in all sampled sections, with all periosteal regions showing active primary bone deposition, indicating that both individuals were still growing at the time of death (Figs 8B and 8G). MMNS VP-6332 exhibits a severe and chronic pathology affecting the cortex, the detailed description of which is beyond the scope of this paper.

**Fig 8.**
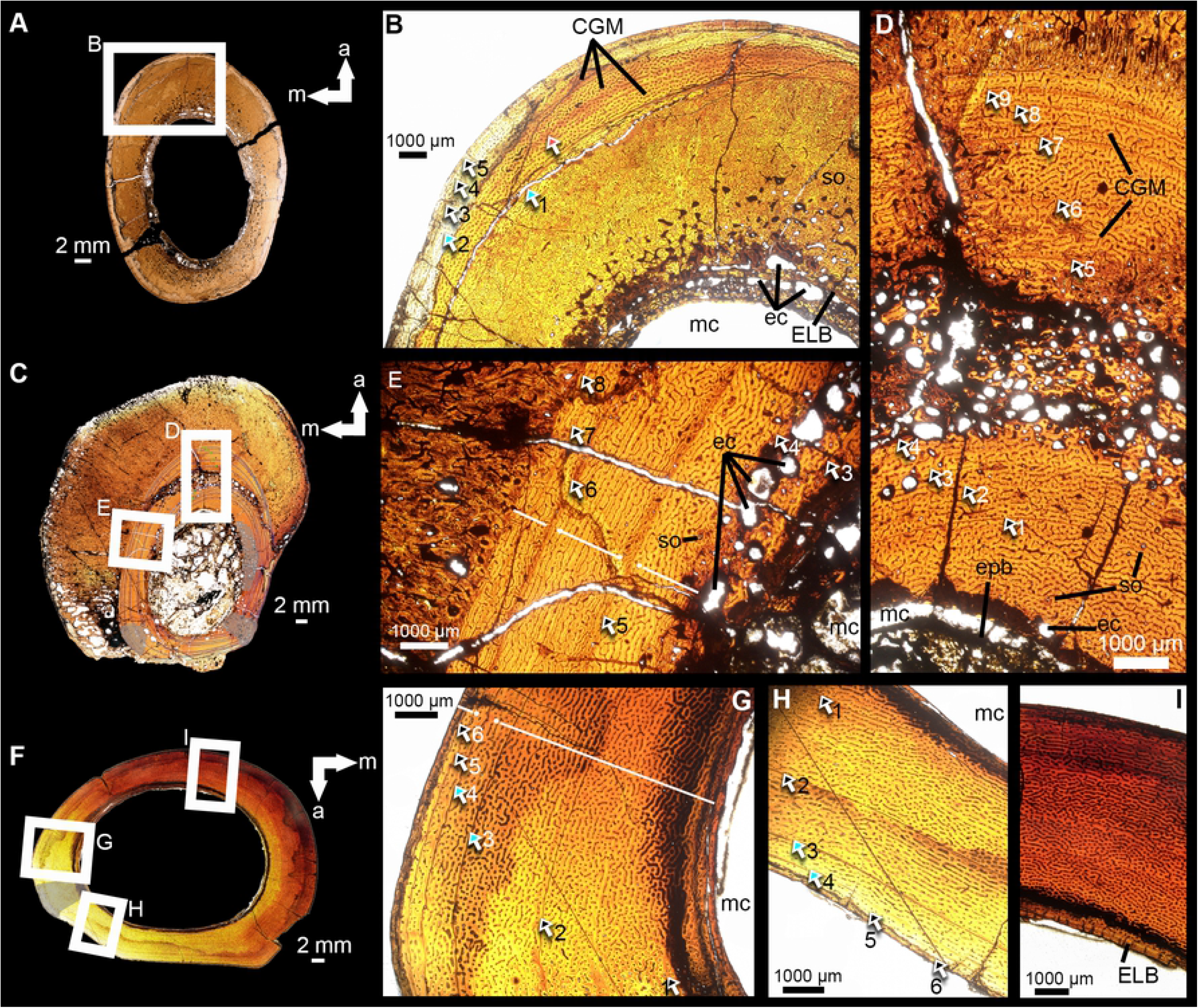
Paleohistological transverse sections of selected Eutaw ornithomimosaur elements. Cross-sections of the proximal-most (**A-B**), and midsection (**C-E**) of the second metatarsal shaft (MMNS VP-6332) of a large-bodied individual, and mid-section (**F-I**) of the tibial shaft (MMNS VP-7649) of medium-bodied individual. Details of the pattern of vascular orientation, LAG counts, cyclical growth marks, and remodeling shown (E-D), and (G-I). (B) the anterior-anteromedial region of image-A, showing primary tissue including lines of arrested growth (LAGs); (E-D) details of the pattern and the orientation of the vascularization of the anterior and the medial regions of the cortex of the midsection. Abbreviations: a, anterior; cgm, cyclical growth mark; elb, endosteal lamellar bone; epb, endosteal pathologic bone; m, medial; mc, medullary cavity; rc, resorptive cavity; so, secondary osteon. LAGs are indicated by green arrowheads in images (B), (D-E), and (G-H). Dashed white line in image (G) indicates the transitional boundary of the zonation of vascularity. Gray-shaded areas in images (C) and (F) indicate the distribution of the secondary osteons in the cortex.

#### The cross-section of the proximal shaft of MMNS VP-6332

MMNS VP-6332 was transversely sectioned just distal to the proximal-most end of the shaft (Fig 8A). The thickness of the cortex ranges from 7.64-11.02 mm and is thickest anteriorly (Fig 8A). A large medullary cavity is lined with well-preserved endosteal lamellar bone (ELB) composed of multiple layers of avascular tissue (Fig 8B). Portions of the ELB are obliterated by endosteal deposition of pathologic bone and resorption cavities (Fig 8A-B). Whereas the anteromedial and medial regions of the outer cortex are composed of primary bone, the inner two-thirds of the entire cortex is extensively remodeled circumferentially by multiple generations of secondary osteons (Haversian bone). Between the ELB and this region of secondary remodeling, numerous similarly-sized erosive cavities extend from the endosteal junction of the cortex centrifugally to the middle cortex, forming a spongy medullary region, except for a few large cavities, concentrated along the perimedullary region (Fig 8B). The overall density of secondary osteons gradually decreases periosteally, so that the primary bone tissue is visible in the outer region. Primary bone tissue consists of a woven-fibered matrix. In the outer cortex, the orientation of the vascularization is predominantly longitudinal and reticular, with a minor component of sub-plexiform vascularization (Fig 8B). At least six cyclical growth marks, including five lines of arrested growths (LAGs) and one annulus, are present in the region of primary tissue between the periosteal extent of secondary remodeling and the periosteal margin. The first two LAGs include a couplet (Fig 8B). Together these comprised seven cyclical growth marks, although other LAGs were likely lost as a result of the extensive remodeling of the inner cortex due to the pathology and medullary expansion (Fig 8A-B). The three LAGs closest to the periosteal surface are tightly packed, superficially similar to the condition of an EFS. However, the matrix remains woven in this region, the tissue is vascularized, and osteocyte lacunae are relatively plump, suggesting that chronic pathology may more likely be the cause of the close spacing in these LAGs.

#### The cross-section of the midshaft of MMNS VP-6332

An additional section of this element was made from the midshaft to capture the longest growth record; however, the pathology is more extensive in this region (Fig 8C). The bone matrix is a predominantly woven-parallel complex (Fig 8C-E). The thickness of the cortex is relatively consistent in the lateral, medial, and posterior regions, and ranges between 4.59-5.64 mm, but the anterior cortex is more than twice this thickness (12.06 mm). The medullary cavity is large (maximum diameter 22.39 mm) relative to the cortex, which is infilled by highly disorganized pathologic bone obliterating most of the primary bone near the endosteal surface (Fig 8C-D). The vascular orientation in the cortex changes centripetally beginning with predominantly reticular vascularization near the endosteal margin, transitioning to a combination of plexiform and laminar vascularization in the middle cortex, and leading to primarily plexiform vascularization towards the periosteal surface (visible medially). This is generally consistent with the vascular patterns reported for other ornithomimid metatarsals [112,113] (Fig 8E). Secondary osteons are rare throughout the cortex, except for three concentrated areas of remodeling with cross-cutting secondary osteons in the lateral, posterolateral, and posteromedial regions (Fig 8C, gray shaded areas). In addition, some narrow zones are infilled with secondary osteons, indicative of repaired cracks in the cortex. Erosion cavities are extensive throughout the cortex and are relatively large (0.08-1.6 mm), substantially larger than those of the proximalmost shaft of MMNS VP-6332. Remnants of the ELB (tightly packed avascular lamellar-fibered bone tissue) are present at the posterolateral margin of the medullary cavity. At least nine LAGs are traceable throughout the cortex of the midsection, representing ten growth cycles (Fig 8C-D). The pattern of LAG spacing is relatively constant in the innermost four LAGs of the inner half of the cortex, with somewhat more distant spacing in the mid-cortex, and a consistent decrease in spacing in the remaining LAGs in the outer cortex towards the periosteal surface (Fig 8D), which is consistent with a pattern often observed in sub-adult theropods [90]. No definitive EFS is visible in the periosteal surface, consistent with the condition of the proximalmost shaft, confirming that an EFS is not present in this individual (Fig 8D-E).

#### The cross-section of the midshaft of MMNS VP-7649

The cortex of the MMNS VP-7649 midshaft is predominantly composed of a woven-fiber matrix dominated by reticular and plexiform vascularization and areas of increased density of radial canals (Fig 8G). There are few to no secondary osteons throughout the cortex except for a localized zone of secondary osteons across an endosteal-periosteal gradient in the anterolateral region (Fig 8F). Unlike the metatarsal MMNS VP-6332, there is no evidence of substantial cortical drift in the tibial section (MMNS VP-7649) via uneven expansion of the medullary cavity (Fig 8F). The thickness of the cortex appears relatively consistent relative to the large medullary cavity, although the medial side is thicker on average. There is zonation in the vascularity of the outer cortex, most notably near the outer region of the lateral cortex beginning with the fourth LAG (Fig 8G). Whereas much of the inner cortex is highly vascularized up to the fourth LAG, the cortex between the fourth LAG and the periosteum is less densely vascularized, as evidenced by the presence of fewer vascular canals (Fig 8G). Six LAGs are present from the innermost endosteal region to the periosteal surface, representing at least seven growth cycles, and the third and fourth LAGs contain a couplet (Fig 8G). LAG spacing is inconsistent throughout the cortex of MMNS VP-7649, but generally decreases substantially, with wide spacing between LAGs in the inner cortex and tighter, decreasing spacing in the outer cortex (Fig 8G-H). There is no definitive presence of the EFS in the periosteum (Fig 8G-I).

## Discussion

Ornithomimosaurs repeatedly evolved gigantic body size during their evolutionary history [14,70,114–116] (Fig 9), although evidence for directional mass evolution, as opposed to stochastic processes, is lacking [117]. Early-diverging ornithomimosaurs from the Early Cretaceous (pre-Albian), such as *Nqwebasaurus thwazi* [93] from Africa, *Hexing qingyi* [103], *Shenzhousaurus orientalis* [104], and *Kinnareemimus khonkaenensis* [105] from Asia, *Pelecanimimus polyodon* [106] from Europe, and *Nedcolbertia justinhofmanni* [107] from North America) were universally small bodied (>12 kg [102; table S1]). During the Albian, ornithomimosaurs generally embarked on a trend of increasing body size, although a mosaic of small, medium, and large bodied species existed (Figs 2 and 9). This is known, in part, based on ornithomimosaur remains from the Cloverly Formation [108] and Arundel Clay [15], *H. okladnikovi* from Mongolia [109] (all small to medium-bodied species), as well as the earliest examples of large bodied species with some taxa, such as *A. fridayi* in North America [14] and *B. grandis* in Asia [70], exceeding 350 kg [37] (S1 Table). By the end of the Cretaceous (Campanian-Maastrichtian), multiple large-bodied species are known to have inhabited Laurasian landmasses, including the deinocheirid *Paraxenisaurus normalensis*, from the Campanian Cerro del Pueblo Formation of Mexico, indeterminate large-bodied ornithomimid materials from the Dinosaur Park Formation, Canada [25,27,116,125,126] and *G. bullatus* (MPC-D 100/11) from the Nemegt Formation of Mongolia [64]. Moreover, by this time ornithomimosaurs had achieved gigantism, as exemplified by *D. mirificus* (MPC-D 100/127), which is estimated to have weighed over 6,000 kg [110] (S1 Table).

**Fig 9.**
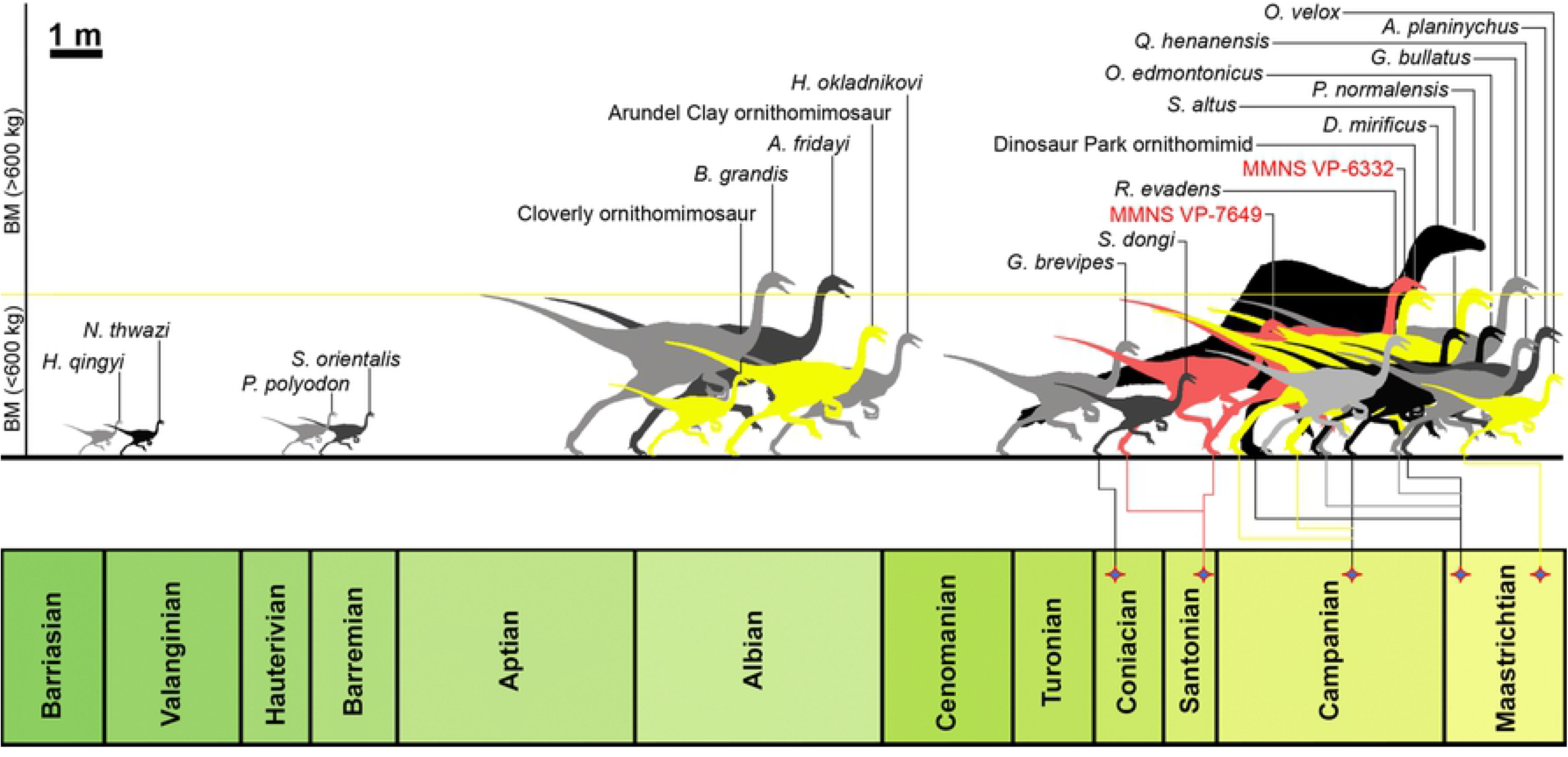
Relative body-size of the Eutaw ornithomimosaurs and geologic age of known ornithomimosaur taxa. Estimated relative body sizes are based on the femoral lengths data obtained from Zanno and Makovicky [117]. All silhouettes are *Ornithomimus*, except for *D. mirificus*. Yellow silhouettes indicate that the relative body mass is estimated from fragmentary elements. Silhouette courtesy Mr. Keiji Terakoshi.

Despite an increase in our understanding of ornithomimosaurian body size evolution during the Cretaceous period, substantial gaps persist. The North American specimen record is poor between the Albian and Campanian [27,125]; similarly, little is known about ornithomimosaur mass evolution between the Turonian/Coniacian and Maastrichtian across Asia. The presence of large-bodied Eutaw ornithomimosaurs from the Santonian of Mississippi fills a gap in the record of body size evolution of ornithomimosaurs in North America and further suggests the presence of coexisting large- and small/medium-bodied forms in the region during this time (Figs 2 and 9).

Using extant-scaling approaches from estimated femur circumferences, we estimate the body mass range of three individuals within our Eutaw sample. The upper and the lower bounds for each of these three individuals are ranged between 607-1025 kg for MMNS VP-6332, 6790-11470 kg for MMNS VP-7119, and 136-230 kg for MMNS VP-7649, when *D. mirificus* is included in the regression. Conversely, the upper and the lower bounds for the same individuals are estimated at 221-374 kg for MMNS VP-6332, 884-1494 kg for MMNS VP-7119, and 105-177 kg for MMNS VP-7649, when *D. mirificus* is excluded from the regressions (S1 Table). This is broadly consistent with the mass ranges of other large-bodied ornithomimosaurs (e.g., *A. fridayi*, ∼380 kg; *B. grandis*, ∼375 kg; *G. bullatus*, ∼400 kg), which, on their maximum end, are intermediate between the masses of these taxa and the largest ornithomimosaur *D. mirificus* (∼6350 kg) (see S1 Table for comparative mass estimates using this and other mass estimation approaches). However, body mass estimation of the smaller individual (MMNS VP-7649) is consistent with the size of the reported late-diverging ornithomimosaur taxa from the Late Cretaceous of North America (S1 Table). It should also be emphasized that the regressions performed to estimate the FCs (and subsequently estimate BM) used different elements, based on the availability of material from MMNS VP-6332 (metatarsal), MMNS VP-7119 (pedal phalanx), and MMNS VP-7649 (tibia), and that each of these element experiences differing allometric effects that can impact the accuracy when predicting values that are substantially larger than the original data used to perform the regression. This may be partially responsible for the particularly large FC and BM estimates in MMNS VP-7119, when compared to the estimates in MMNS VP-7649 and MMNS VP-6332.

Among large-bodied ornithomimosaurs, growth data based on bone histology is limited to *B. grandis* [70]. Based on a cross-section of the fibular shaft, the holotype specimen of *B. grandis* belongs to an individual that was at least 13–14 years old and actively growing, although approaching somatic maturity, at the time of death. Our osteohistological data suggests that the second metatarsal MMNS VP-6332 belongs to an individual minimally 10 years old and still growing at death (Fig 3C-D). Decreasing LAG spacing toward the periosteal surface, and the absence of an EFS, suggests a subadult individual of similar relative growth stage as *B. grandis*.

Cullen and colleagues [113] performed multi-element osteohistological analysis on the hind limb elements of the Dry Island ornithomimid specimens to assess intra- and interskeletal variations, such as the difference in the number of LAGs within different elements of the same individual. They documented that in that sample, the fibula generally preserved one more LAG than the femur, tibia, or metatarsal, although this pattern was not always true in histological samplings among theropods more broadly [90,113,127]. Using this relationship to qualitatively approximate the number of missing LAGs and ontogenetic age in years, we find that the estimated age and size are grossly similar between these two taxa, with the likelihood that the Eutaw taxon represented by MMNS VP-6332 would be substantially larger at skeletal maturity, even if it stopped growing at a younger age. For comparison, the holotype of *B. grandis* is estimated at ∼13–14 years of age and ∼375 kg (based on extant-scaling approaches using femoral circumference; see S1 Table), and the Eutaw ornithomimosaur represented by the second metatarsal (MMNS VP-6332) is estimated as at least 11 years of age (estimating for missing LAGs and metatarsal-fibular growth mark record differences) and weighing ∼816 kg based on extant-scaling approaches using femoral circumference (see S1-S2 Tables for other mass estimates).

In contrast, the tibia (MMNS VP-7649) from our Eutaw ornithomimosaur assemblage likely represents a distinct and smaller-bodied taxon. This specimen preserves seven growth cycles (Fig 8G), compared to the 10 preserved by the second metatarsal (MMNS VP-6332) (Fig 8D), yet belongs to an individual with a body mass estimated to be ∼20-50% the body mass of MMNS VP-6332 (depending on inclusion/exclusion of *D. mirificus* in predictive FC and BM dataset for Eutaw specimens; Fig 9 and S1 Fig; Table 1 and S1 Table). The combination of osteohistological and femoral circumference/mass data from our sample strongly suggests the co-occurrence of at least two distinct ornithomimosaur taxa--one small and one large-bodied--in the Santonian of southern Appalachia. Alternatively, it is possible that the smaller tibia represents an earlier ontogenetic stage of the larger taxon, but we find this inconsistent with the current data given that the tibia (MMNS VP-7649) preserves closer spacing of outer LAGs and decreasing vascular complexity (Fig 8G), suggesting that the animal was approaching maturity, despite recording three to four fewer growth cycles than the much larger metatarsal of MMNS VP-6332. While differences in average LAG spacing have been documented between ornithomimosaur metatarsals and major long-bones (e.g., femora, tibiae; [76,98]), such intra-skeletal growth record variations are unlikely to account for the substantial differences in growth mark count and mass estimates.

The coexistence of two ornithomimosaur genera is relatively common in Cretaceous ecosystems of Laurasia [15,118,119,128,129]. Typically, co-occurrences include taxa of similar body size (small or medium bodied) [118,119]. However, co-occurrences of both small/medium and large-bodied ornithomimosaur genera are rare. Few examples include the Late Cretaceous Nemegt Formation (early Maastrichtian) of Mongolia, which preserves the medium-bodied taxa *Anserimimus planinychus* and *G. bullatus* and large-bodied taxon *D. mirificus* [115,129,130], and the Dinosaur Park Formation (mid to late Campanian) of Canada, including the medium-bodied taxa *O. velox* and *S. altus*, and potentially also an unnamed large ornithomimosaur [126].

The presence of large-bodied (>350 kg) ornithomimosaur taxa from the late Early Cretaceous through Maastrichtian of North America including *A. fridayi* from the Aptian/Albian of Arkansas, a proximal tibia (SMU 76809) of large ornithomimosaur from the Cenomanian Lewisville Formation of Texas [22], Ornithomimosauria indet. materials described herein from the Santonian of Mississippi, *P. normalensis* from the Campanian of Mexico, and a large unnamed ornithomimid (LACM 47520) from the late Maastrichtian Hell Creek Formation of Montana, together with Asian representatives from the Early Cretaceous (*B. grandis*) and the Late Cretaceous (*D. mirificus* and *G. bullatus*) of China and Mongolia, indicate that large-bodied ornithomimosaurs had achieved a near pan-Laurasian distribution by the post-Aptian Cretaceous (post-Aptian).

## Conclusions

To date, the assemblage of ornithomimosaur materials from the Santonian Eutaw Formation are some of the best-preserved theropod materials known from Upper Cretaceous sediments of Appalachia. Specimens described herein add essential new information to the poorly known mid-Cretaceous interval from the Appalachian landmass by filling a critical gap in the spatiotemporal and biodiversity records of ornithomimosaurs in North America, documenting the youngest occurrence of ornithomimosaurs in Appalachia (during an interval of faunal isolation), the only definitive Santonian record of ornithomimosaurs on the North American continent, and one of the largest ornithomimosaurian species known globally. This record, when combined with the previously described Arundel Clay ornithomimosaurs, *A. fridayi*, and a partial tibia of a large unnamed ornithomimosaur from the Lewisville Formation of Texas, confirm that ornithomimosaur dinosaurs were present in Appalachia throughout the early Late Cretaceous.

Previous studies on the dissociated ornithomimosaur specimens have demonstrated that manual and/or pedal elements of ornithomimosaurs are important source of taxonomically informative anatomical information and can be diagnostic for Ornithomimosauria [30,65,126]. This is supported by several studies describing ornithomimosaurians on the basis of solely manual or pedal elements, for example *A. tugrikinensis*, *A. planinychus*, *A. fridayi*, *P. normalensis*, and *T. packardensis* [14,62,92,116,131]. Despite this, the likelihood of more than one co-occurring taxon in the Eutaw assemblage, coupled with a lack of association between elements, and the presence of pathologies, prevents us from confidently assigning the Eutaw specimens to finer taxonomic levels, such as a species, in this time. Nonetheless, the presence of two ornithomimosaur taxa in the Santonian Eutaw assemblage, based on the combined size and growth data presented herein, is consistent with Campanian-Maastrichtian assemblages of the North America and Asia [115,129], suggesting that multiple species of ornithomimosaurian theropods likely cohabited within Laurasian ecosystems throughout the latter half of the Late Cretaceous. Due to gaps in the fossil record, it is currently unclear if this pattern is related to the evolution of large body size (>350 kg) among some ornithomimosaurians initiating in the Aptian/Albian (e.g., *B. grandis*, *A. fridayi*). Evolution of large body size in select taxa would be expected to correlate with a niche shift and such a trend could explain the presence of multiple ornithomimosaur taxa in Late Cretaceous ecosystems, as well as the lack of evidence for directional evolution in body mass via the co-occurrence of multiple clades of different sized ornithomimosaurians through geologic time. It is interesting that some of the largest ornithomimosaurs known from North America stem from an interval of high sea-level and reduced range area (Santonian). Robust paleobiogeographic analyses that would elucidate patterns of ornithomimosaurian dispersal across Laurasian landmasses up to the Turonian (specifically between North America and East Asia during the mid-Cretaceous, a.k.a EKLInE, [122]) must await the discovery of additional and more complete material. The same is true regarding population isolation in Laramidia and Appalachia on and its effect on evolutionary trajectories in body-size.

## Acknowledgments

We gratefully acknowledge those who collected and donated the materials used in this project to public repositories including Eric Loftis, Jason Robinson, Wilkie Collins, Joe Gibson, Neal Larson, David Troskey, Chuck Ciampaglio, Pete Larson, and James Lamb. We also express our gratitude to the landowners Keith and Meredith Hester and Richard Fleming, as well as the City of Columbus, for unrestricted access to the fossil exposures along Luxapallila Creek. We also thank the following people: Lisa Herzog and Aurore Canoville (Paleontological Research Laboratory, NC Museum of Natural Sciences) for assistance with the procedures of molding/casting and histological samplings, respectively; Greg Funston (University of Edinburgh), Yoshitsugu Kobayashi, and Philip Currie (University of Alberta) for sharing their specimen photos and useful for the comparisons; Adrian Smith (Evolutionary Biology & Behavior Laboratory, Natural Research Center, NC Museum of Natural Sciences) for providing facilities necessary for microscopic investigations; and Rebecca Hunt (BLM, Canyon Country District, Utah), Thomas Holtz (University of Maryland), Terry Gates (NC State University), Ryuji Takasaki (Okayama University of Sciences), Junki Yoshida (Fukushima Museum), and Gregory S. Paul for helpful discussions on the earlier versions of the manuscript and during the annual meeting of the Society of Vertebrate Paleontology. We are grateful to Keiji Terakoshi for permission to use the silhouette image of *Ornithomimus*. Funding was provided by the NC Museum of Natural Sciences and the Mississippi Museum of Natural Science.

## Supporting information captions

**S1 Fig.** Graphs of linear regression analysis of the Eutaw ornithomimosaur elements. Femoral circumference versus (**A1-A2**), pedal phalanx IV-2 lengths, (**B1-B2**), metatarsal II lengths, and (**C1-C2**) tibial lengths. Note that the graphs of A1, B1, and C1 show when *D. mirificus* is included in the analysis and A2, B2, and C2 show when *D. mirificus* is excluded from the analysis.

**S2 Fig.** Comparison of the astragali of theropod dinosaurs. (**A**), *T. rex* (MOR 1125); (**B**), *D. aquilunguis* (ANSP 9995); (**C**), *A. montgomeriensis* (RMM 6670); (**D**), *F. utahensis* (UMNH VP 12364); (**E**), *Anzu* sp. (NCSM 33801); (**F**), *T. sampsoni* (UMNH VP 19479); (**G**), *Q. henanensis* (left, HGM 41HIII-0106); (**H**), *A. tugrikinensis* (MPC-D 100/130); (**I**), Bissekty taxon (ZIN PH 144/16). Abbreviations: alr, anterolateral ridge; ap, ascending process; asc, articular surface for the calcaneum; asf, articular surface for the fibula; bap, base of the ascending process; cal, calcaneum; fos, median fossa; hg, horizontal groove; icb, intercondylar bridge; jun, junction; lc, lateral condyle; mc, medial condyle; n, notch; slf, laterally flared articular surface of the base; sluf, laterally unflared articular surface of the base; Note that all astragali refer to left except for the right side of *F. utahensis* and is reversed. Images are adapted and modified from (B), Brusatte et al. [102]; (C), Carr et al. [11]; (F), Zanno et al. [57]; (G), Xu et al. [60]; (I), Sues and Averianov [105].

**S3 Fig.** Comparison of second metatarsals of tyrannosauroids. (**A**), *A. montgomeriensis* (RMM 6670); (**B**), *A. atokensi*s (NCSM 14345); (**C**), *A. sarcophagus* (AMNH 5432); (**D**), *T. rex* (FMNH PR 2081). Note that all images refer to the right metatarsals except for the left side of *A. sarcophagus*. Images adapted and modified from (A), Carr et al. [11]; (C-D), Brochu [63]. Not to scale.

**S4 Fig.** Comparison of the distal halves of third metatarsals of ornithomimosaurs. (**A**), *O. velox* (YPM 542); (**B**), *D. brevitertius* (UA 16182); (**C**), *R. evadens* (ROM 1790); (**D**), *A. tugrikinensis* (MPC-D 100/130); (**E**), Large Gansu ornithomimid metatarsal (IVPP 12756); (**F**), *Q. henanensis* (HGM 41HIII-0106); (**G**), *A. fridayi* (UAM 74-16); (**H**), *B. grandis* (FRDC-GS GJ 06). Abbreviations: acdn, distally mediolaterally not widened articular caput; acdw, distally mediolaterally widened articular caput; Note that all images refer to left third metatarsals except for the right side of large Gansu ornithomimid and *Arkansaurus fridayi*. Images are adapted and modified from (C), McFeeters et al. [74]; (E), Shapiro et al. [65]; (F), Xu et al. [60]; (G), Hunt and Quinn [14]; (H), Makovicky et al. [70]. Not to scale.

**S5 Fig.** Comparison of the distal halves of third metatarsals of theropod dinosaurs. (**A**), *C. pergracilis* (=*M. canadensis*), (CMN 8538); (**B**), *E. rarus* (MPC-D 102/6); (**C**), *A. montgomeriensis* (RMM 6670); (**D**), *T. sampsoni* (UMNH VP 19479); (**E**), *S. inequalis* (=*L. mcmasterae*) (TMP 1992.036.0575). Abbreviations: elp, extensor ligament pit; ics, intercondylar sulcus; lc, lateral condyle; lt, “lateral tab,”; mc, medial condyle. Note that all images refer to the right metatarsals except for the left metatarsal of *T. sampsoni*. Images adapted and modified from (B), Currie et al. [133]; (C), Carr et al. [11]; (D), Zanno et al. [57]; (E), van der Reest and Currie [89]. Image of the left metatarsus of *T. sampsoni* is mirrored. Not to scale.

**S6 Fig.** Comparison of first pedal phalanges of third digit of selected theropod dinosaurs. (**A**), *C. pergracilis* (=*M. canadensis*) (CMN 8538); (**B**), *A. wyliei* (CMN 78000); (**C**), juvenile *T. bataar* (MPC-D 107/7); (**D**), *G. libratus* (FMNH PR 2211); (**E**), *Alectrosaurus* sp. (MPC-D 100/51); (**F**), *A. sarcophagus* (CMN 11315); (**G**), *T. sampsoni* (UMNH VP-19479); (**H**), *F. utahensis* (); (**I**), *G. erlianensis* (LH V0011); (**J**), *A. atokensis* (NCSM 14345); (**K**), *A. fragilis* (YPM 1930); (**L**), *T. rex* (FMNH PR 2081). Abbreviations: elp, extensor ligament pit. Note that A, D, F, J, and L are referred to the right and B, C, E, G-I, and K are referred to the left pedal phalanges. Images adapted and modified from (G), Zanno et al. [57]; (H), Zanno [59]; (K), Madsen [94], and A and L are mirrored. Not to scale.

**S7 Fig.** The preserved vertebrae of the Eutaw ornithomimosaurs. (**A**), the anterior dorsal (MMNS VP-6120); (**B**), the posterior dorsal (MMNS VP-113); (**C**), the posterior caudal centra. (A1-C1), left lateral; (A2-C2), right lateral; (A3-C3), dorsal; (A4-C4), ventral; (A5-C5), anterior; and (A6-C6), posterior views. Abbreviations: am, angled margin of the articular surface; ar, angular ridge; dep, depression; gr, groove; lr, longitudinal groove; na, neural arch; nc, neural canal; ncs, neurocentral suture; ns, neural spine; poz, postzygapophysis; prz, prezygapophysis; vk, ventral keel. Scale bars equal to 3 cm for A_1_-A_4_ – C_1_-C_4_ and 2 cm for A_5_-A_6_ – C_5_-C_6_.

**S8 Fig.** Comparison of the anterior (A) and the posterior (B) dorsal vertebrae of selected theropod dinosaurs. **A1, C1-D1**, the fourth and **A2,C2-D2,** the eleventh dorsal vertebrae; (A1-A2), *D. antirrhopus* in left lateral views; (B), *S. meekerorum* in right (B1) and left (B2) lateral views; (C1-C2), *A. riocoloradense* in left lateral views; (D1-D2), *T. rex* (FMNH PR2081) in right (D1) and left (D2) lateral views. Abbreviations: ncs, neurocentral suture; pa, parapophysis; pl, pleurocoel; pnf, pneumatic fossa. Images (A1-A2) and (C1) are reversed. Images adapted and modified from (A1-A2), Ostrom [111]; (B1-B2), Zanno and Makovicky [134]; (C1-C2), Sereno et al. [135]; (D1-D2), Brochu [63]. Not to scale.

**S9 Fig.** Comparison of the posterior caudal vertebrae of selected theropod dinosaurs. (**A**), *D. aquilunguis*; (**B**), *A. montgomeriensis*; (**C**), *M. roseae*; (**D**), *Z. salleei*; (**E**), *T. rex*; (**F**), *G. libratus*; (**G**), *T. euotica*; (**H**), Utah tyrannosaurid. Abbreviations: na, neural arch; ns, neural spine; poz, postzygapophysis; prz, prezygapophysis. Images adapted and modified from (A), Brusatte et al. [102]; (B), Carr et al. [11]; (C), Coria and Currie, [103]; (D), Choiniere et al. [136]; (E), Brochu [63]; (F), Lambe [87]; (G), Brusatte et al. [72]; (H), Loewen et al. [137]. Not to scale.

**S10 Fig.** Manual phalanges of the Eutaw ornithomimosaurs. (**A**), the right first phalanx of digit 3 (PhIII-1); (**B**), the ungual phalanx (I-1?). (A1, B3), anterior; (A2, B4), posterior; (A3, B1), lateral; (A4, B2), medial; (A5, B5), proximal; and (A6), distal views. Abbreviations: clf, collateral ligament fossa; conc, concavity; dep, depression; elp, extensor ligament pit; ft, flexor tubercle; glm, ginglymoid articular surface; ics, intercondylar sulcus; lc, lateral condyle; lgr, lateral groove; mc, medial condyle; mr, medial ridge; p, process; t, tubercle.

**S11 Fig.** Partial tibia of the medium-bodied Eutaw ornithomimosaur. **(A)**, anterior; **(B)**, posterior; **(C)**, lateral; **(D)**, medial; **(E)**, proximal; **(F)**, distal views. Interpretive illustration of *Q. henanensis* (HGM 41HIII-0106) shows the approximate location of the preserved portion of the midshaft. Abbreviations: a, anterior; bcc, a base of the cnemial crest; cfb, contact surface for the fibula; fc, fibular crest; l, lateral; m, medial; p, posterior.

**S12 Fig.** Comparison of the tibiae of selected theropod dinosaurs. (**A**), *G. bullatus* (MPC-D 100/11); (**B**), *Q. henanensis* (HGM 41HIII-0106); (**C**), *B. grandis* (FRDC-GS GJ 06); (**D**), *F.utahensis* (UMNH VP 12362); (**E**), *M. intrepidus* (NCSM 33392); (**F**), *D. aquilunguis* (ANSP 9995); (**G**), *A. montgomeriensis* (RMM 6670); (**H**), *T. rex* (FMNH PR 2081). Abbreviations: agr, a groove of the articular facet for the fibula; fc, fibular crest; nf, nutrient foramen. A, B, D, F, and H are referred to left and C, E, and G are referred to the right tibiae. Images adapted and modified from (A), Osmólska et al. [64]; (B), Xu et al. [60]; (C), Makovicky et al. [70]; (D), Zanno [59]; (E), Zanno et al. [86]; (F), Brusatte et al. [102]; (G), Carr et al. [11]; (H), Brochu [63]. All images are seen from lateral view except for (A) and (G), which are in posterior view. Not to scale.

**S1 Table.** Measurement comparisons of the select pedal elements of ornithomimosaurs. Note that a single asterisk (*) indicates the measurements including *Deinocheirus mirificus*, and a double asterisk (**) indicate the measurements excluding *Deinocheirus mirificus*.

| Species | Specimen # | Pedal phalanx IV-2 Length | Metatarsal II Length | Tibia Length | Femur circumference (FC) |
| --- | --- | --- | --- | --- | --- |
| *Anserimimus planinychus* | MPC-D100/300 | 17 | 270 | 450 | 134 |
| *Archaeornithomimus asiaticus* | AMNH 6570 | - | 274 | 301 | 46 |
| *Deinocheirus mirificus* | MPC-D 100/127 | 64 | 510 | 1140 | 560 |
| *Dromiceiomimus brevitertius* | AMNH 5201 | 20 | - | 438 | 103 |
| *Dromiceiomimus brevitertius* | ROM 797 | 30 | 253 | 470 | 99 |
| *Dromiceiomimus brevitertius* | ROM 852 | 29 | 325 | 520 | 128 |
| *Dromiceiomimus brevitertius* | UALVP 16182 | - | 305 | 473 | 131 |
| *Gallimimus bullatus* | MPC-D100/10 | 13 | 144 | 218 | 54 |
| *Gallimimus bullatus* | MPC-D100/11 | 43 | 480 | 695 | 195 |
| *Gallimimus bullatus* | MPC-D100/12 | - | 330 | 508 | 160 |
| *Gallimimus bullatus* | MPC-D100/52 | 11 | 256 | 400 | 128 |
| *Gallimimus bullatus* | MPC-D100/121 | 23 | 278 | 450 | 130 |
| *Gallimimus bullatus* | MPC-D100/123 | 31 | - | 695 | 180 |
| *Gallimimus bullatus* | MPC-D100/138 | 29 | 458 | 685 | 165 |
| *Gallimimus bullatus* | ZPal MgD-I/1 | 17 | 264 | 384 | 109 |
| *Gallimimus bullatus* | ZPal MgD-I/32 | 22 | - | - | 133 |
| *Gallimimus bullatus* | ZPal MgD-I/94 | 19 | 205 | 292 | 76 |
| *Ornithomimidae* | MPC-D100/8 | 29 | - | - | 115 |
| *Ornithomimidae* | MPC-D100/136 | 13 | - | - | 100 |
| *Garudimimus brevipes* | MPC-D100/13 | 35 | 195 | 388 | 107 |
| *Ornithomimus edmontonicus* | ROM 851 | 29 | 265 | 475 | 110 |
| *Ornithomimus edmontonicus* | TMP 95.110.1 | 22 | 297 | 465 | 125 |
| *Struthiomimus altus* | AMNH 5257 | 31 | 342 | 555 | 160 |
| *Struthiomimus altus* | AMNH 5375 | - | 357 | 408 | 141 |
| *Struthiomimus altus* | UCMZ (VP) 1980.1 | 27 | - | 556 | 136 |
| *Rativates evadens* | ROM 1790 | 20 | 277 | 421 | 111 |
| **Eutaw ornithomimosaur** | **MMNS VP-6332** | - | 435 | - | 252.09* |
|  |  | - | 435 | - | 174.68** |
| **Eutaw ornithomimosaur** | **MMNS VP-7649** | - | - | 506 | 146.45* |
|  |  | - | - | 506 | 133.13** |
| **Eutaw ornithomimosaur** | **MMNS VP-7119** | 92 | - | - | 606.87* |
|  |  | 92 | - | - | 289.09** |

**S2 Table.**
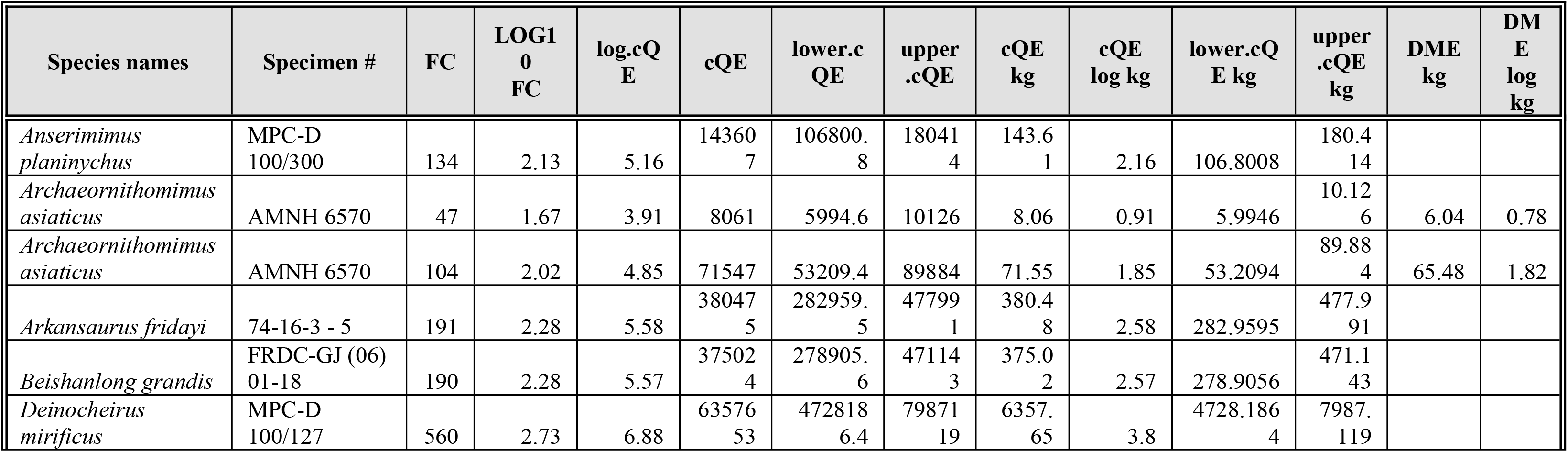
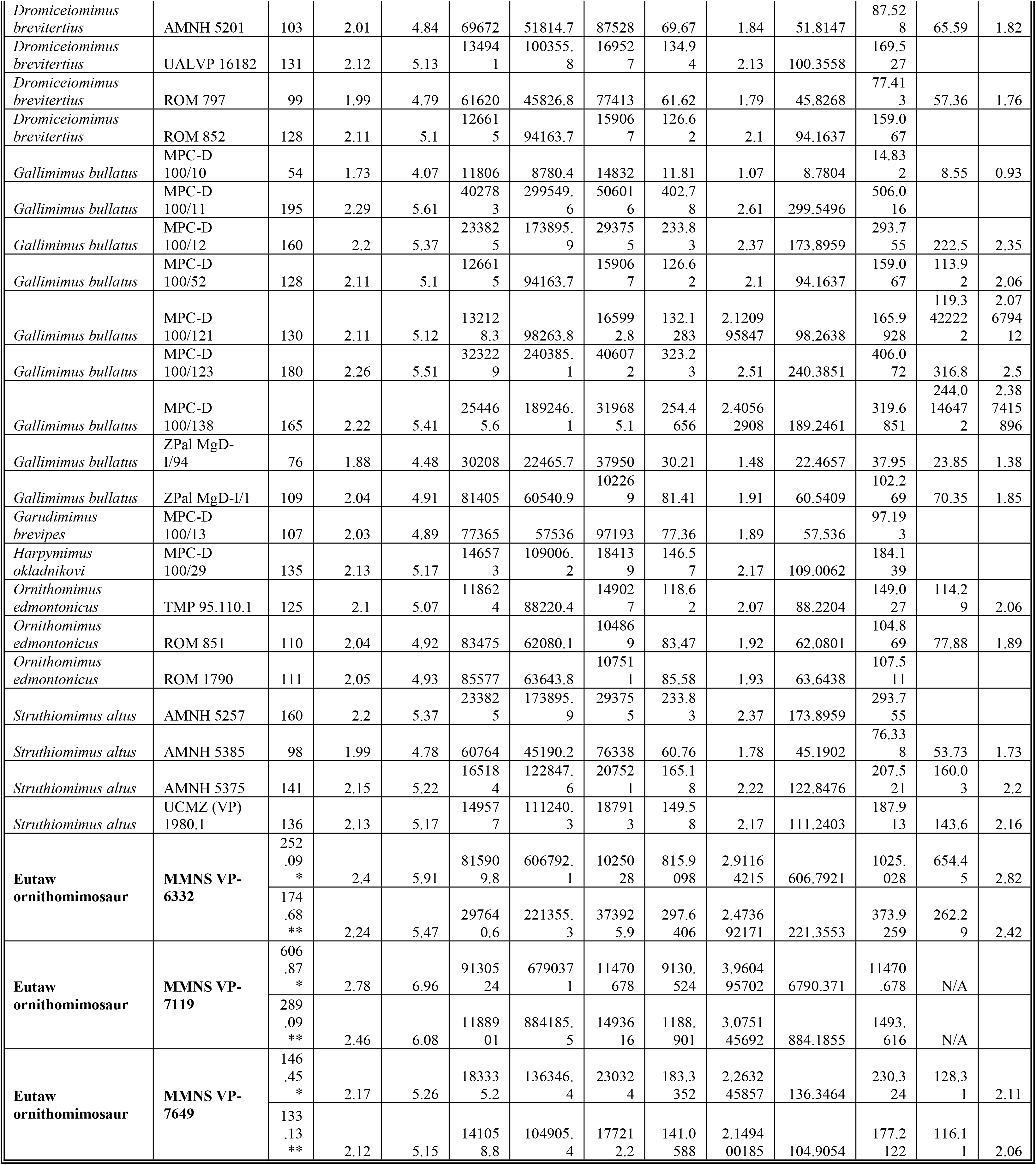
Body mass estimates for ornithomimosaurs. Note that a single asterisk (*) indicates the values when D. mirificus is included, and double asterisk (**) indicate the values when *Deinocheirus mirificus* is excluded in the analysis. Abbreviations: (cQE), the corrected quadrupedal values for biped; (DME), developmental mass extrapolation; (FC), femoral circumference.

